# The coding capacity of SARS-CoV-2

**DOI:** 10.1101/2020.05.07.082909

**Authors:** Yaara Finkel, Orel Mizrahi, Aharon Nachshon, Shira Weingarten-Gabbay, David Morgenstern, Yfat Yahalom-Ronen, Hadas Tamir, Hagit Achdout, Dana Stein, Ofir Israeli, Adi Beth-Din, Sharon Melamed, Shay Weiss, Tomer Israely, Nir Paran, Michal Schwartz, Noam Stern-Ginossar

**Author notes:** These authors contributed equally to this work (Yaara Finkel, Orel Mizrahi).

## Abstract

Severe acute respiratory syndrome coronavirus 2 (SARS-CoV-2) is the cause of the ongoing Coronavirus disease 19 (COVID-19) pandemic ^1,2^. In order to understand SARS-CoV-2 pathogenicity and antigenic potential, and to develop diagnostic and therapeutic tools, it is essential to portray the full repertoire of its expressed proteins. The SARS-CoV-2 coding capacity map is currently based on computational predictions and relies on homology to other coronaviruses. Since coronaviruses differ in their protein array, especially in the variety of accessory proteins, it is crucial to characterize the specific collection of SARS-CoV-2 proteins in an unbiased and open-ended manner. Utilizing a suite of ribosome profiling techniques ^3–8^, we present a high-resolution map of the SARS-CoV-2 coding regions, allowing us to accurately quantify the expression of canonical viral open reading frames (ORF)s and to identify 23 novel unannotated viral translated ORFs. These ORFs include upstream ORFs (uORFs) that are likely playing a regulatory role, several in-frame internal ORFs lying within existing ORFs, resulting in N-terminally truncated products, as well as internal out-of-frame ORFs, which generate novel polypeptides. We further show that viral mRNAs are not translated more efficiently than host mRNAs; rather, virus translation dominates host translation due to high levels of viral transcripts. Overall, our work reveals the full coding capacity of SARS-CoV-2 genome, providing a rich resource, which will form the basis of future functional studies and diagnostic efforts.

## Main

SARS-CoV-2 is an enveloped virus consisting of a positive-sense, single-stranded RNA genome of ~30 kb and shows characteristic features of other coronaviruses. Upon cell entry, two overlapping ORFs are translated from the positive strand genomic RNA, ORF1a and ORF1b. The translation of ORF1b is mediated by a −1 frameshift that allows translation to continue into ORF1b enabling the generation of continuous polypeptides which are cleaved into a total of 16 nonstructural proteins (NSPs) ^9–11^. In addition, the viral RNA-dependent RNA polymerase (RdRP) uses the viral genome to produce negative-strand RNA intermediates which serve as templates for the synthesis of positive-strand genomic RNA and of subgenomic RNAs ^9–11^. The subgenomic RNAs contain a common 5’ leader fused to different segments from the 3’ end of the viral genome, and contain a 5’-cap structure and a 3’ poly(A) tail ^12,13^. These unique fusions occur during negative-strand synthesis at 6-7 nt core sequences called transcription-regulating sequences (TRS)s that are located at the 3’ end of the leader sequence as well as preceding each viral gene. The different subgenomic RNAs encode 4 conserved structural proteins-spike protein (S), envelope protein (E), membrane protein (M), nucleocapsid protein (N)- and several accessory proteins. Based on sequence similarity to other beta coronaviruses and specifically to SARS-CoV, current annotation of SARS-CoV-2 includes predictions of six accessory proteins (3a, 6, 7a, 7b, 8, and 10, NCBI Reference Sequence: NC_045512.2), but not all of these ORFs have been experimentally and reproducibly confirmed in this virus^14,15^.

To capture the full SARS-CoV-2 coding capacity, we initially applied a suite of ribosome profiling approaches to Vero E6 cells infected at MOI=0.2 with SARS-CoV-2 (BetaCoV/Germany/BavPat1/2020 EPI_ISL_406862) for 5 or 24 hours (Figure 1A). At 24 hours post infection the vast majority of cells were infected but cells were still intact (Figure S1). For each time point we mapped genome-wide translation events by preparing three different ribosome-profiling libraries (Ribo-seq), each one in two biological replicates. Two Ribo-seq libraries facilitate mapping of translation initiation sites, by treating cells with lactimidomycin (LTM) or harringtonine (Harr), two drugs with distinct mechanisms that inhibit translation initiation by preventing 80S ribosomes formed at translation initiation sites from elongating. These treatments lead to strong accumulation of ribosomes precisely at the sites of translation initiation and depletion of ribosomes over the body of the message (Figure 1A and ^4,6^). The third Ribo-seq library was prepared from cells treated with the translation elongation inhibitor cycloheximide (CHX), and gives a snap-shot of actively translating ribosomes across the body of the translated ORF (Figure 1A). These three complementary approaches provide a powerful tool for the unbiased mapping of translated ORFs. In parallel, RNA-sequencing (RNA-seq) was applied to map viral transcripts. When analyzing the different Ribo-seq libraries across coding regions in cellular genes, the expected distinct profiles are observed in both replicates, confirming the overall quality of the libraries. Ribosome footprints displayed a strong peak at the translation initiation site, which, as expected, is more pronounced in the Harr and LTM libraries, while the CHX library also exhibited a distribution of ribosomes across the entire coding region up to the stop codon, and its mapped footprints were enriched in fragments that align to the translated frame (Figure 1B, Figure S2 and Figure S3). As expected, the RNA-seq reads were uniformly distributed across coding and non-coding regions (Figure 1B and Figure S2). The footprint profiles of viral coding sequences at 5 hours post infection (hpi) fit the expected profile of translation, similar to the profile of cellular genes, both at the meta gene level and at the level of individual genes (Figure 1C, Figure S4 and Figure S5). In addition, the footprint densities at 5hpi were highly reproducible between our biological replicates both at the gene level and in single codon resolution on the viral genome (Figure S6). Intriguingly, the footprint profile over the viral genome at 24 hpi, did not fit the expected profile of translating ribosomes and were generally not affected by Harr or LTM treatments (Figure S4). To further examine the quality of footprint measurements we applied a fragment length organization similarity score (FLOSS) that measures the magnitude of disagreement between the footprint distribution on a given transcript and the footprint distribution on canonical CDSs ^16^. At 5 hpi protected fragments from SARS-CoV-2 transcripts showed the expected size distribution (Figure S7A) and scored well in these matrices and did not differ from well-expressed human transcripts (Figure 1D). However, reads from 24 hpi could be clearly distinguished from cellular annotated coding sequences (Figure 1E and Figure S7B). We conclude that the footprint data from 5hpi constitutes robust and reproducible ribosome footprint information but that the majority of viral protected fragments at 24 hpi are likely not generated by ribosome protection and may reflect additional interactions that occur on the viral genome at late time points in infection.

**Figure 1.**
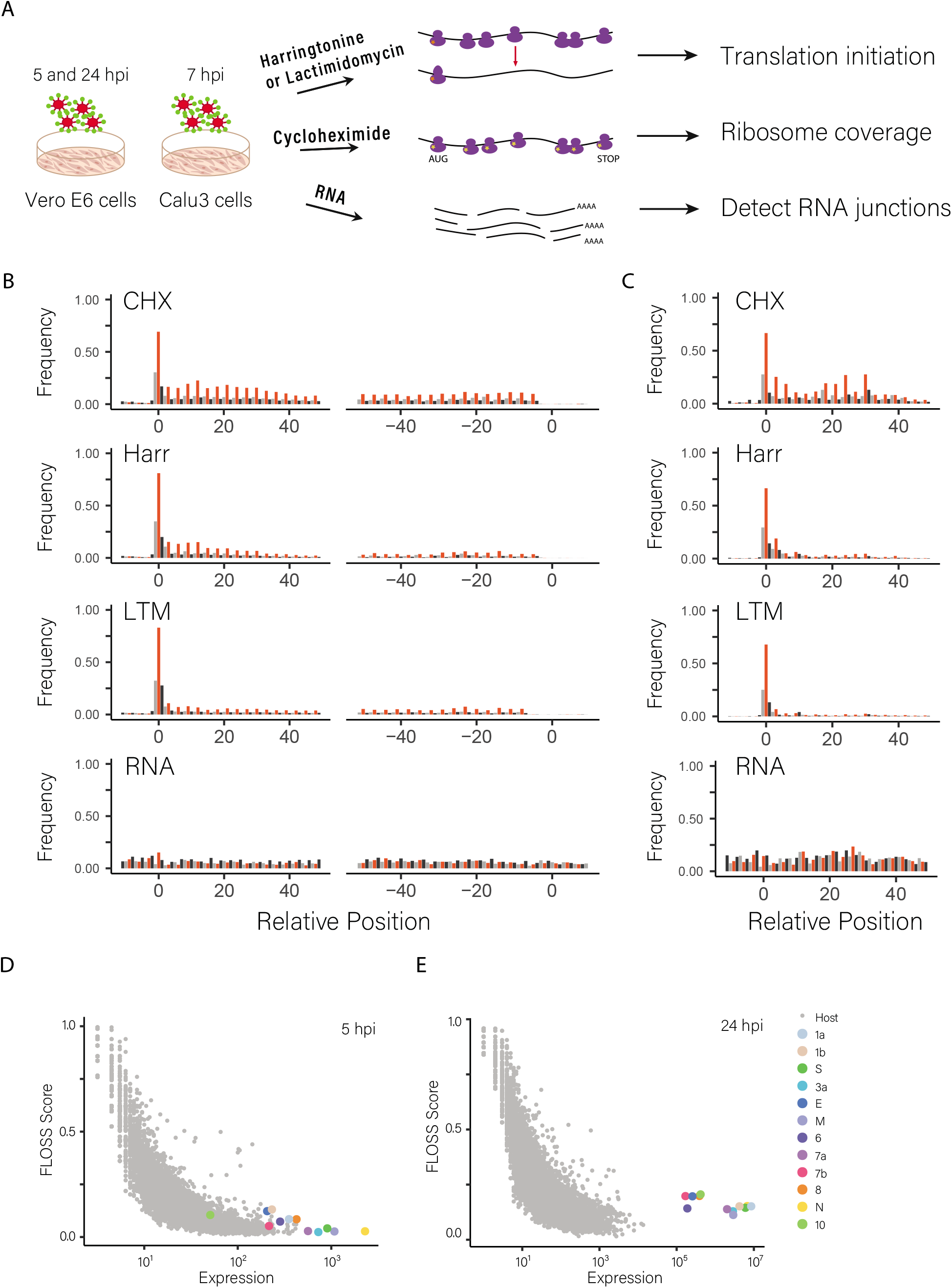
Ribosome profiling of SARS-CoV-2 infected cells. **(A)** Vero E6 and Calu3 cells infected with SARS-CoV-2 were harvested at 5, 24 (Vero E6) and 7 (Calu3) hours post infection (hpi) for RNA-seq, and for ribosome profiling using lactimidomycin (LTM) and Harringtonine (Harr) treatments for mapping translation initiation or cycloheximide (CHX) treatment to map overall translation. **(B)** Metagene analysis of read densities at the 5’ and the 3’ regions of cellular protein coding genes as measured by the different ribosome profiling approaches and RNA-seq at 5 hpi (one of two replicates is presented). The X axis shows the nucleotide position relative to the start or the stop codons. The ribosome densities are shown with different colors indicating the three frames relative to the main ORF (red, frame 0; black, frame +1; grey, frame +2). **(C)** Metagene analysis of the 5’ region, as described in B, for viral coding genes at 5 hpi **(D and E)** Fragment length organization similarity score (FLOSS) analysis for cellular coding regions and for SARS-CoV-2 canonical ORFs at 5 hpi **(D)** and 24 hpi **(E)**.

A global view of RNA and CHX footprint reads mapping to the viral genome at 5hpi, demonstrate an overall 5’ to 3’ increase in coverage (Figure 2A). RNA levels are essentially constant across ORFs 1a and 1b, and then steadily increases towards the 3’, reflecting the cumulative abundance of these sequences due to the nested transcription of subgenomic RNAs (Figure 2A). Increased coverage is also seen at the 5’ UTR reflecting the presence of the 5’ leader sequence in all subgenomic RNAs as well as the genomic RNA. Reduction in footprint density between ORF1a and ORF1b reflects the proportion of ribosomes that terminate at the ORF1a stop codon instead of frameshifting into ORF1b (Figure S8). By dividing the footprint density in ORF1b by the density in ORF1a we estimate frameshift efficiency is 57% +/− 12%. This value is comparable to the frameshift efficiency measured based on ribosome profiling of mouse hepatitis virus (MHV, 48%-75%)^7^. On the molecular level this 57% frameshifting rate indicates NSP1-NSP11 are expressed 1.8 +/− 0.4 times more than NSP12-NSP16 and this ratio likely relates to the stoichiometry needed to generate SARS-CoV-2 nonstructural macromolecular complexes ^17^. Similarly to what was seen in MHV and avian infectious bronchitis virus (IBV) ^7,8^, we failed to see noticeable ribosome pausing before or at the frameshift site, but we identified several potential pausing sites within ORF1a and in ORF1b that were reproducible between replicates (Figure S8), however these will require further characterization.

**Figure 2.**
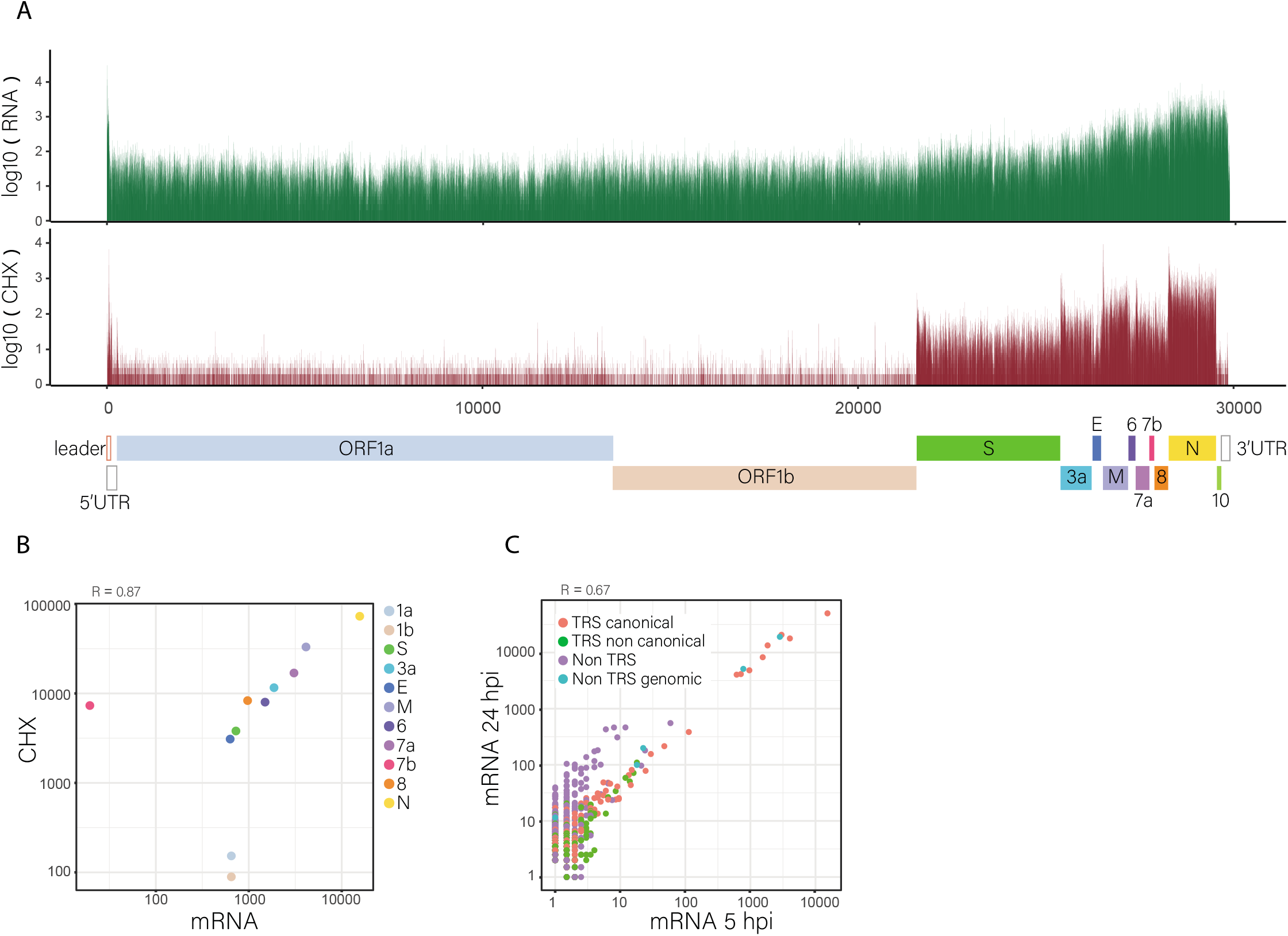
Expression level of canonical viral genes. **(A)** RNA-Seq (green) and Ribo-Seq CHX (red) read densities at 5 hpi on the SARS-CoV-2 genome. Read densities are plotted on a log scale to cover the wide range in expression across the genome. The lower panel presents SARS-CoV-2 genome organization with the canonical viral ORFs **(B)** Relative abundance of the different viral transcripts relative to the ribosome densities of each SARS-CoV-2 canonical ORF at 5 hpi. Transcript abundance were estimated by counting the reads that span the junctions of the corresponding RNA or for ORF1a and ORF1b the genomic RNA abundance, normalized to junction count. ORF10 is not presented as no junctions designated for its subgenomic RNA were detected. Spearman’s R is presented. **(C)** Scatter plot presenting the abundance of viral reads that span canonical leader dependent junctions (red), non-canonical leader dependent junctions (green), non-canonical leader independent junctions (purple) and genomic deletions (cyan) at 5 and 24 hpi. Pearson’s R on log transformed values is presented.

Besides ORF1a and ORF1b, all other canonical viral ORFs are translated from subgenomic RNAs. We therefore examined whether the levels of viral gene translation correlate with the levels of the corresponding subgenomic RNAs. Since raw RNA-seq densities represent the cumulative sum of genomic and all subgenomic RNAs, we calculated transcript abundance using two approaches: deconvolution of RNA densities, in which RNA expression of each ORF is calculated by subtracting the RNA read density of cumulative densities upstream to the ORF region; and relative abundances of RNA reads spanning leader-body junctions of each of the canonical subgenomic RNAs. ORF6, ORF7b and ORF10 obtained negative values in the RNA deconvolution, probably due to their short length and relative weaker expression, which make them more sensitive to inaccuracies related to library preparation biases. For ORF10 we also did not detect reads spanning leader-body junctions. For all other canonical ORFs there was high correlation between these two approaches (Pearson’s R = 0.897, Figure S9), and in both approaches the N transcript was the most abundant transcript, in agreement with recent studies ^15,18^. We next compared footprint densities to RNA abundance as calculated by junction abundances for the subgenomic RNA or deconvolution of genomic RNA in the case of ORF1a and ORF1b (Figure 2B). For the majority of viral ORFs, transcript abundance correlated almost perfectly with footprint densities, indicating these viral ORFs are translated in similar efficiencies (probably due to their almost identical 5’UTRs), however three ORFs were outliers. The translation efficiency of ORF1a and ORF1b was significantly lower. This can stem from unique features in their 5’UTR (discussed below) or from under estimation of their true translation efficiency as some of the full-length RNA molecules may serve as template for replication or packaging and are hence not part of the translated mRNA pool. The third outlier is ORF7b for which we identified very few body-leader junctions but exhibited relatively high translation. A probable explanation is that translation of ORF7b arises from ribosome leaky scanning of the ORF7a transcript, as was suggested in SARS-CoV ^19^.

Recently, many transcripts derived from non-canonical junctions were identified for SARS-CoV-2, some of which were abundant and were suggested to affect the viral coding potential ^15,18^. These non-canonical junctions contain either the leader combined with 3’ fragments at unexpected sites in the middle of ORFs (leader-dependent noncanonical junction) or fusion between sequences that do not have similarity to the leader (leader-independent junction). We estimated the frequency of these non-canonical junctions in our RNA libraries. We indeed identified many non-canonical junctions and obtained excellent agreement between our RNA-seq replicates for both canonical and non-canonical junctions, demonstrating these junctions are often generated and mostly do not correspond to random amplification artifacts (Figure S10A, S10B and Table S1). The abundance of junction-spanning reads between our data and the data of Kim et al. ^18^, that was generated from RNA harvested from Vero cells at 24 hpi, showed significant correlation (Pearson’s R = 0.816 for 24hpi, Figure S10C and S10D), illustrating many of these are reproducible between experimental systems. However, 111 out of the 361 most abundant leader independent junctions that were mapped by Kim et al., were not supported by any read in our data, illustrating there are also substantial variations. In addition we identified 5 abundant leader independent junctions that were not expressed based on Kim et al. ^18^ (Table S2). We noticed three of these junctions represent short in-frame deletions in the spike protein (5aa, 7aa and 10aa long) that overlap deletions that were recently described by other groups ^15,20,21^, in which the furin-like cleavage site is deleted (Figure S11). The re-occurrence of the same genomic deletion strongly supports the conclusion that this deletion is being selected for during passage in Vero cells. In order to examine if any additional non-canonical junctions are derived from genomic deletions we sequenced the genomic RNA of the virus we used in our infections. Indeed, 50% of the genomic RNA contained a 5-10 aa deletion of the furin-like cleavage site in the spike protein. In addition, we identified an 8aa deletion in ORF-E in 2.3% of the genomic RNA, which was also observed in our RNA measurements (Table S2 and Figure 2C). When we compared the frequency of junctions between 5h and 24h time points, the leader dependent junctions (both canonical and non-canonical) and the genomic deletions correlated well but the leader independent junctions were specifically increased at 24 hpi (Figure 2C). Recent kinetic measurements show viral particles already bud out of infected Vero cells at 8 hpi ^21^. Therefore, this time-dependent increase in non-canonical RNA junctions indicates that the leader independent RNA junctions are likely associated with genomic replication. Overall, this data shows a small part of the leader-independent junctions represent genomic deletions that are likely selected for during cell culture passages and a larger subset of leader-independent junctions probably rises during genome replication and therefore less likely to lead to changes in viral transcripts.

Our ribosome profiling approach facilitates unbiased assessment of the full range of SARS-CoV-2 translation products. Examination of SARS-CoV-2 translation as reflected by the diverse ribosome footprint libraries, revealed several unannotated translated ORFs. We detected in-frame internal ORFs lying within existing ORFs, resulting in N-terminally truncated product. These included relatively long truncated versions of canonical ORFs, such as the one found in ORF6 (Figure 3A and Figure S12A), or very short truncated ORFs that likely serve an upstream ORF (uORF), like truncated ORF7a that might regulate ORF7b translation (Figure 3B, Figure S12B and Figure S12C). We also detected internal out-of-frame translations, that would yield novel polypeptides, such as ORFs within ORF-3a (41aa and 33 aa, Figure 3C and Figure S12D) and within ORF-S (39aa, Figure 3D and Figure S12E) or short ORFs that likely serve as uORFs (Figure 3E and Figure S12F). Additionally, we observed a 13 amino acid extended ORF-M, in addition to the canonical ORF-M, which is predicted to start at the near cognate codon AUA (Figure 3F and Figure S12G and Figure S12H).

**Figure 3.**
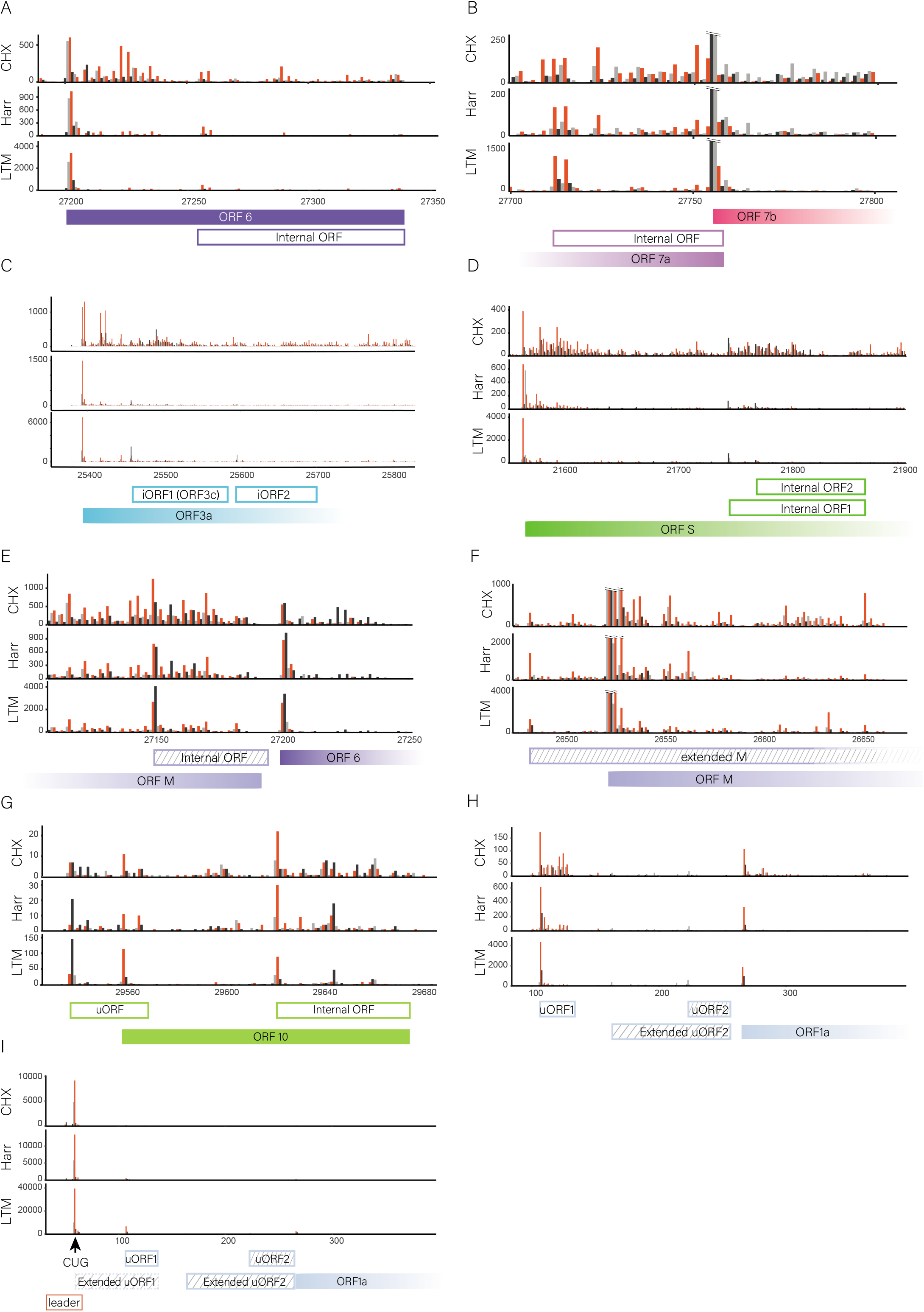
Ribosome densities reveal novel viral coding regions. **(A-I)** Ribosome density profiles of CHX, Harr and LTM samples at 5 hpi. Densities are shown with different colors indicating the three frames relative to the main ORF in each figure (red, frame 0; black, frame +1; grey, frame +2). One out of two replicates is presented. Filled and open rectangles indicate the canonical and novel ORFs, respectively. ORFs starting in a near cognate codon are labeled with stripes. ORFs that stretch beyond the range of the plot are shown as fading out rectangles. **(A)** In frame internal ORF within ORF6 generating a truncated product, **(B)** In frame internal initiation within ORF7a (reads marking ORF-7b initiation were cut to fit the scale indicated with black lines), **(C)** Out of frame internal initiations within ORF-3a, **(D)** Out of frame internal initiations within ORF-S, **(E)** Out of frame internal initiation within ORF-M (the predominant junction for ORF6 is upstream of this iORF, outside the range displayed, and is not shown in the figure), **(F)** an extended version of ORF-M (reads marking ORF-M initiation were cut to fit the scale indicated with black lines), **(G)** uORF that overlap ORF10 initiation and in frame internal initiation generating truncated ORF10 product **(H)** two uORFs embedded in ORF1a 5’UTR **(I)** non canonical CUG initiation upstream of the TRS leader.

The presence of the annotated ORF10 was recently put into question as almost no subgenomic reads were found for its corresponding transcript ^18,22^. Although we also did not detect subgenomic RNA designated for ORF10 translation (Table S1), the ribosome footprint densities indicate translation signal in ORF10 initiation (Figure 3G and Figure S12I). Interestingly, we detected two putative ORFs, an upstream out of frame ORF that overlaps ORF10 initiation and an in-frame internal initiation that leads to a truncated ORF10 product. Further research is needed to delineate how ORFs in this region are translated and whether they have any functional roles.

Finally, we detected four distinct initiation sites at SARS-CoV-2 5’UTR. Three of these encode for uORFs that are located just upstream of ORF1a; the first initiating at an AUG (uORF1) and the other two at a near cognate codons (uORF2 and extended uORF2, Figure 3H and Figure S12J). These uORFs are in line with findings in other coronaviruses ^7,23^. The fourth site is the most prominent peak in the ribosome profiling densities on the SARS-CoV-2 genome and is located on a CUG codon at position 59, just 10 nucleotides upstream the TRS-leader (Figure 3I and Figure S12K). The reads mapped to this site have a tight length distribution characteristic of ribosome protected fragments (Figure S13A). Due to its location upstream of the TRS-leader, footprints mapping to this site can potentially derive from any of the subgenomic as well as the genomic RNAs. Therefore, to view this initiation in its context, we aligned the footprints to the genomic RNA or to the most abundant subgenomic N transcript. On the genome and on ORF-N transcript this initiation results in translation of uORFs, which on the genome will generate an extension of uORF1 (Figure S13B and Figure S13C). In both transcripts, the occupancy at the CUG is higher than the downstream translation signal, implying this peak might reflect ribosomal pausing. Interestingly, ribosome pauses located just upstream of the TRS-leader were also identified in MHV and IBV genomes ^7,8^. To assess the distribution of footprints at this initiation on the different viral transcripts, viral transcripts were divided into three groups based on their sequence similarity downstream of the leader-junction site (to allow unique footprint alignment, Figure S13D). Interestingly, significantly more footprints were mapped to the group that includes the genomic RNA and the subgenomic E and M transcripts, than would be expected from their relative RNA abundance (Figure S13E). When only footprints that allow unique mapping to genomic RNA or subgenomic M and E transcripts are used (sizes 31-33bp to discriminate M from genome or E transcript, and sizes 32-33bp to discriminate E from the genome) a strong enrichment of footprints that originate from the genome is observed (Figure S13F). This footprint enrichment to genomic RNA suggests ribosome pausing might be more prominent on the genome or that ribosomes engage with genomic RNA differently than with subgenomic transcripts. The proximity of this pause to the leader-TRS, which seem to be conserved in MHV and IBV ^7,8^, together with the relative enrichment to the viral genome raises the possibility that a ribosome at this position might affect discontinuous transcription either by sterically blocking the TRS-L site or by affecting RNA secondary structure. In addition, ribosomes initiating at the CUG have the potential to generate uORFs or ORF extensions in the different sub-genomic transcripts (Table S3)

To systematically define the SARS-CoV-2 translated ORFs we used PRICE and ORF-RATER, two computational methods that rely on a combination of translation features such as LTM and Harr induced peak at the translation initiation site, heightened footprints density and 3-nt periodicity to predict novel translated ORFs from ribosome profiling measurements ^24,25^. After application of a minimal expression cutoff and manual curation on the predictions, these classifiers identified 25 ORFs, these included 10 out of the 11 canonical translation initiations and 15 novel viral ORFs. In addition, ORF-RATER identified three putative ORFs that originate from the CUG initiation and extend to the sub-genomic transcripts of S, M and ORF-6a (Table S3). The majority (85%) of the classifier identified ORFs where independently identified in each of the biological replicate (Table S4 and Figure S14). Visual inspection of the ribosome profiling data suggested additional 8 putative novel ORFs, some of which are presented above (Figure 3A, 3B, 3G and Table S4). Overall, we identified 23 putative ORFs, on top of the 12 canonical viral ORFs that are currently annotated in NCBI Reference Sequence and 3 additional potential ORFs that stem from the CUG initiation upstream of the leader.

To confirm the robustness and relevance of these annotations we extended these experiments to human cells. We first examined the infection efficiency of several human cell lines that were used to study SARS-CoV-2 infection; epithelial lung cancer cell lines, Calu3 and A549, and epithelial colorectal adenocarcinoma cell line, Caco2. Infection of Calu3 was most efficient and infection in the presence of trypsin increased infection efficiency by at least 2-fold (Figure S15). We next infected Calu3 with a SARS-CoV-2 that was isolated from an independent source (BavPat1/2020 Ref-SKU: 026V-03883) and the integrity of the virus that we used for infection was confirmed by sequencing (confirming the virus sequence was intact and there were no abundant genomic deletions). Cells were harvested at 7hpi using the same set of ribosome profiling techniques, each one in two biological replicates and in parallel RNA was harvested for RNA-seq. The different Ribo-seq libraries showed the expected distinct profiles in both replicates, confirming the overall quality of these libraries (Figure S16). We next examined the translation of the new viral ORFs we have annotated using PRICE and ORF-RATER classifiers as well as manual curation. Of the 23 novel ORFs we identified as being translated in Vero cells all showed clear evidence of translation also in Calu3 infected cells, 16 were annotated by PRICE and ORF-RATER (three of which are ORFs that were added manually based on the Vero cells data) and ORF-RATER identified again the same three ORFs that originate from the CUG initiation upstream the leader (Figure S17, Table S3 and Table S4). LTM-induced ribosome accumulation at the canonical and predicted initiation sites were highly reproducible between biological replicates as well as between Calu3 and Vero cells, illustrating the robustness of the translation initiation predictions (Figure S18A-C). Furthermore, ribosome-protected footprints displayed a 3-nt periodicity that was in phase with the predicted start site, in both Vero and Calu3 cells providing further evidence for the active translation of the predicted ORFs (Figure S19). We conclude 23 novel ORFs are reproducibly translated from SARS-CoV-2 independently of the host cell and the exact viral origin and additional ORFs may be translated from the CUG initiation located upstream of the TRS-leader.

Ribosome density also allows accurate quantification of viral protein production. We first quantified the relative expression levels of canonical viral ORFs. Since many of the ORFs we identified overlap canonical ORFs, the quantification was based on the non-overlapping regions. We found that ORF-N is expressed at the highest level in both Vero and Calu3 cells followed by the rest of the viral ORFs with some differences in the relative expression between the two cell types (Figure 4A). To quantify the expression of out-of-frame internal ORFs we computed the contribution of the internal ORF to the frame periodicity signal relative to the expected contribution of the main ORF. For in-frame internal ORF quantification, we subtracted the coverage of the main ORF in the non-overlapping region. We also utilized ORF-RATER, which uses a regression strategy to calculate relative expression of overlapping ORFs, resulting in largely similar estimates of viral ORF translation levels (Figure S20A and S20B). These measurements show that many of the novel ORFs we annotated are expressed in levels that are comparable to the canonical ORFs (Figure 4B, Figure S20C and Table S4). Furthermore, the relative expression of viral proteins seems to be mostly independent of the host cell origin (Figure 4C).

**Figure 4.**
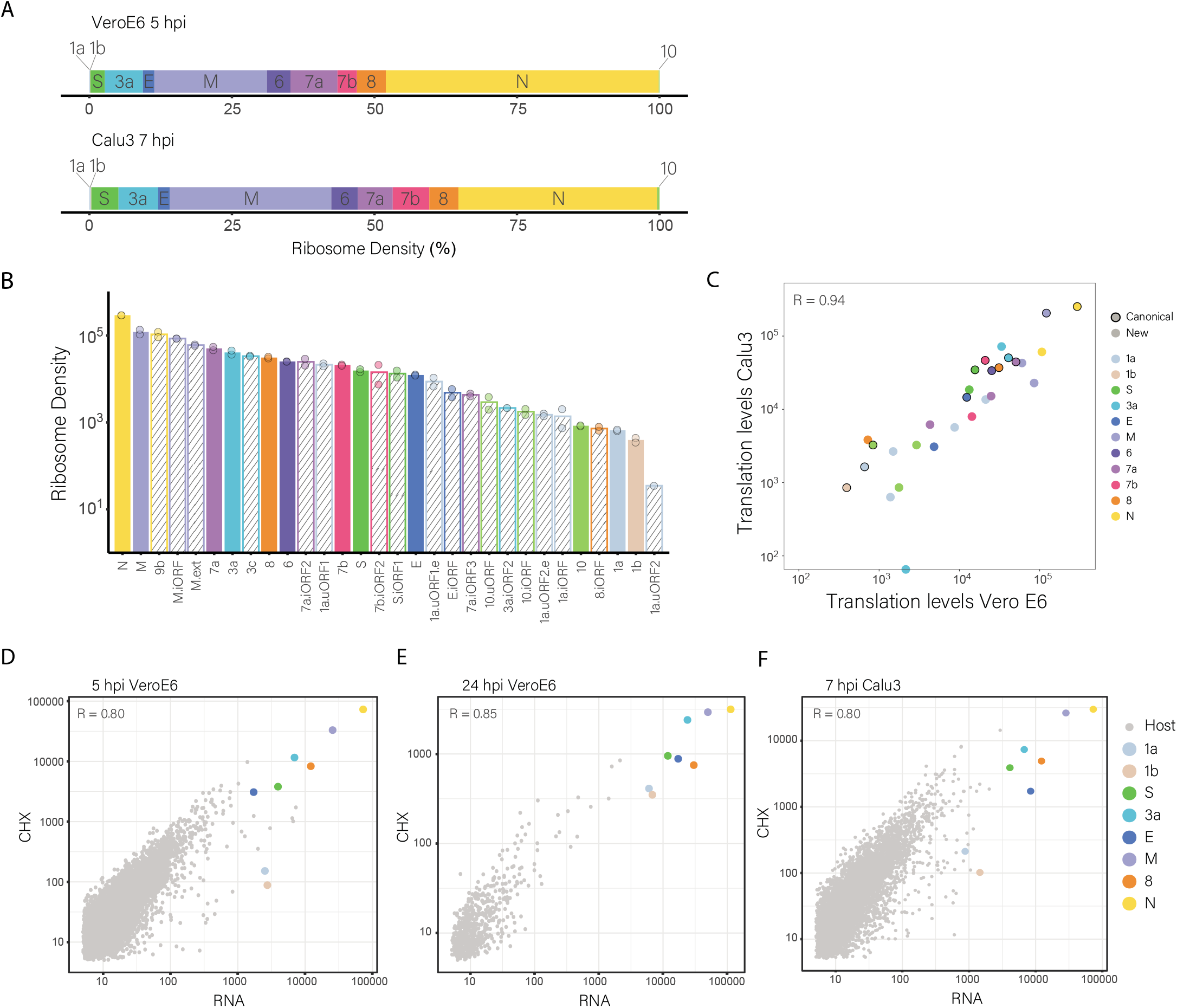
Translation of host and viral genes. **(A)** Relative translation levels of viral coding genes were estimated by counting the ribosome densities on each ORF considering only non-overlapping regions. Data is presented from VeroE6 5hpi and Calu3 7hpi. ORFs are ordered based on their genomic location. **(B)** Viral ORF translation levels as calculated from ribosome densities of infected Vero E6 cells. Data is plotted on a log scale to cover the wide range in expression. Solid fill represents canonical ORFs, and striped fill represents novel ORFs that were annotated. Values were normalized to ORF length and sequencing depth. Points represent the values from the two replicates, except in the case of 3a.iORF2 and 1a.uORF2 where there was a missing value in one of the replicates. **(C)** Scatter plot showing the expression of viral ORFs in infected Vero E6 cells and infected Calu3 cells. Points representing canonical ORFs are outlined in black. Spearman’s R is presented. **(D-F)** Relative transcript abundance versus ribosome densities for each host and viral ORF at 5 hpi **(D)** and 24 hpi **(E)** in Vero E6 cells and at 7hpi in Calu3 cells **(F)**. Transcript abundance was estimated by counting the reads that span the corresponding junction (only the most abundant viral transcripts were counted) and footprint densities were calculated from the CHX sample. For ribo-seq viral reads from 24 hpi, only reads that were mapped to junctions were used to avoid non-ribosome footprints. Pearson’s R on log transformed values is presented for each sample.

Of the novel ORFs we identified 13 are very short (≤ 20 codons) or located in the 5’UTR of the genomic RNA and therefore likely play a regulatory role. Four ORFs are extensions or truncations of canonical ORFs (M, 6, 7a and 10). We examined the properties of the six out-of-frame internal ORFs (iORF)s that are longer than 20aa; one of these is ORF9b and its truncated version (Figure S21A, 97aa and 90aa). ORF9b appears in UniProt annotations and was detected by Bojkova et al. ^14^ in proteomic measurements, together with our translation measurements this indicates it is a bona fide SARS-CoV-2 protein. In addition we detected an iORF laying at the 5’ of ORF-S and its truncated version (Figure 3D, 39 aa and 31 aa), two iORFs within ORF3a (Figure 3C, 41aa and 33 aa). Mining of recent proteomic measurements of SARS-CoV-2 infected cells ^14,15^ did not detect peptides that originate from the out-of-frame ORFs we annotated, likely due to challenges in detecting trypsin-digested products from short coding regions ^24^. Indeed, two canonical SARS-CoV-2 proteins, ORF7b (43aa) and ORF-E (75aa) were also not detected by mass-spectrometry ^14,15,26,27^, and our ribosome profiling data is the first to show these SARS-CoV-2 proteins are indeed expressed.

Using TMHMM, we found S.iORF1 as well as 3a.iORF1 are predicted to contain a transmembrane domain (Figure S22A and S22B). Additionally, using SignalP we found a predicted signal peptide in 3a.iORF2 (Figure S22C). Analysis of the conservation of these out-of-frame iORFs in SARS-CoV and in related viruses (Sarbecoviruses) revealed 3a.iORF1 is highly conserved in Sarbecoviruses (Table S6). This ORF was also identified by three independent comparative genomic studies that demonstrate this ORF has a significant purifying selection signature, implying it is a functional polypeptide^28–30^. In combination with our expression measurements, these findings indicate this internal ORF is a novel and likely functional transmembrane protein, conserved throughout sarbecoviruses and as was suggested by Jungreis et al.^30^ should be named ORF3c.

The second iORF overlapping ORF3a and the iORF overlapping S are not conserved in most other sarbecoviruses (Table S6 and ^29^). The expression of the second iORF overlapping ORF-3a is low (Figure S20) and probably originate from ribosomes that failed to initiate at ORF3a and ORF3c. An extended version of this ORF was pulled-down^31^ and was shown to elicit a strong antibody response ^32^ but we find mainly translation of the short version (Figure S21A). The internal S-ORF is situated just downstream of the ORF-S AUG, suggesting ribosomes might initiate translation via leaky scanning. This region in the S-protein shows extremely-rapid evolution^30^ but in the SARS-CoV-2 isolates that have been sequenced its coding capacity is not impaired. Future work will have to delineate if this ORF, which is highly expressed (Figure 4B and Figure S20), represents a functional protein. Importantly, translated ORFs that do not act as functional polypeptides could still be an important part of the immunological repertoire of the virus as MHC class I bound peptides are generated at higher efficiency from rapidly degraded polypeptides ^33^.

Finally, although we identified two internal out-of-frame ORFs within ORF3a, we did not detect translation of SARS-CoV ORF3b homologue, which contains a premature stop codon in SARS-CoV-2 (Figure S21A). In addition, we did not find evidence of translation of ORF14, which appears in some SARS-CoV-2 annotations ^23^ (Figure S21B).

Translation of viral proteins relies on the cellular translation machinery, and coronaviruses, like many other viruses, are known to cause host shutoff ^34^. In order to quantitatively evaluate if SARS-CoV-2 skews the translation machinery to preferentially translate viral transcripts, we compared the ratio of footprints to mRNAs for virus and host CDSs at 5 hpi and 24 hpi in Vero cells and at 7hpi in Calu3 cells. Since at 24 hpi our ribosome densities were masked by a contaminant signal which do not originate from ribosome protection, for samples from this time point we used the footprints that were mapped to subgenomic RNA junctions (and therefore reflect bona fide transcripts) to estimate the true ribosome densities. In all samples the virus translation efficiencies fall within the low range of most of the host genes (Figure 4D-4F and Figure S23A-C), indicating that viral transcripts are likely not preferentially translated during SARS-CoV-2 infection. Instead, during infection viral transcripts take over the mRNA pool, probably through massive transcription coupled to host induced RNA degradation ^35,36^.

In summary, in this study we delineate the translation landscape of SARS-CoV-2. Comprehensive mapping of the expressed ORFs is a prerequisite for the functional investigation of viral proteins and for deciphering viral-host interactions. An in-depth analysis of the ribosome profiling experiments revealed a highly complex landscape of translation products, including translation of 23 novel viral ORFs and illuminating the relative production of all canonical viral proteins. The new ORFs we have identified may serve as novel accessory proteins or as regulatory units controlling the balanced production of different viral proteins. Studies on the functional significance and antigenic potential of these ORFs will deepen our understanding of SARS-CoV-2 and of coronaviruses in general. Overall, our work reveals the coding capacity of SARS-CoV-2 genome and highlights novel features, providing a rich resource for future functional studies.

## Supporting information

Predicted ORFs bed file

## Tables legend

**Table S1**. Junctions sites detected from junction spanning reads.

This table lists junction sites that were identified by Kim et al. with more than 100 reads and were also detected in our RNA reads.

The genomic coordinates in the “5’ site” and “3’ site” point to the 3’-most and the 5’-most nucleotides that survive the recombination event, respectively. “Gap” is the size of the deletion. “Leader” true value indicates the junction is TRS-leader dependent. “canonical” true value indicates the junction supports the expression of a canonical ORF, “Kim_count” is the number of the junction-spanning reads that support the recombination event identified by Kim et al. “ORF” the name of an ORF that shares the start codon position with the recombination product based on Kim et al. “mrna_05hr_l”, “mrna_05hr_2”, “mrna_24hr_l” and “mrna_24hr_2” the number of the junction-spanning reads that support the recombination event in each of our RNA samples based on STAR-aligner. “fp_chx_05hr_l”, “fp_chx_05hr_2”, “fp_chx_24hr_l” and “fp_chx_24hr_2” the number of the junction-spanning reads that support the recombination event in each of our footprints CHX samples. “sum_fp” the sum of all footprints counts. “sum_mRNA” the sum of all RNA counts. “star_sum” the sum of number of the junction-spanning reads in all samples

**Table S2**. Junctions sites uniquely detected in our samples.

This table lists junction sites that were identified in our RNA samples with more than 50 reads but were low or unidentified by Kim et al.

The genomic coordinates in the “5’ site” and “3’ site” point to the 3’-most and the 5’-most nucleotides that survive the recombination event, respectively. “Gap” is the size of the deletion. “Leader” true value indicates the junction is TRS-leader dependent. “canonical” true value indicates the junction supports the expression of a canonical ORF, “Kim_count” is the number of the junction-spanning reads that support the recombination event identified by Kim et al. “ORF” the name of an ORF that shares the start codon position with the recombination product based on Kim et al. “mrna_05hr_1”, “mrna_05hr_2”, “mrna_24hr_1” and “mrna_24hr_2” the number of the junction-spanning reads that support the recombination event in each of our RNA samples based on STAR-aligner. “fp_chx_05hr_1”, “fp_chx_05hr_2”, “fp_chx_24hr_1” and “fp_chx_24hr_2” the number of the junction-spanning reads that support the recombination event in each of our footprints CHX samples. “sum_fp” the sum of all footprints counts. “sum_mRNA” the sum of all RNA counts. “star_sum” the sum of number of the junction-spanning reads in all samples.

**Table S3**. Potential junction spanning SARS-CoV-2 ORFs that can be translated from the CUG initiation upstream the TRS-leader.

This table lists junction spanning SARS-CoV-2 ORFs that can be translated from the CUG initiation upstream the TRS-leader at each of the sub-genomic RNAs. “Name” for each ORF, “description”, “supported by PRICE Vero” whether the ORF was predicted by PRICE from the Vero E6 data,” supported by ORF-RATER Vero” whether the ORF was predicted by ORF-RATER from the Vero E6 data, “supported by PRICE Calu3” and “supported by ORF-RATER Calu3” whether the ORF was predicted by PRICE or ORF-RATER from the Calu3 data, the “start position” and “end position” in SAS-CoV-2 genome, the nature of the “start codon”, “size(aa)”, “sequence” and “junction position (start, end)”.

**Table S4**. Novel SARS-CoV-2 ORFs that have been identified in our study.

This table lists all the SARS-CoV-2 translated ORFs identified in this study. “Name” for each ORF, “description”, “supported by PRICE Vero” whether the ORF was predicted by PRICE from the Vero E6 data,” supported by ORF-RATER Vero” whether the ORF was predicted by ORF-RATER from the Vero E6 data, “PRICE replicate detection Vero” and “ORF-RATER replicate detection Vero” whether the ORF was detected by PRICE or ORF-RATER using only one replicate if the data, replicate 1 (rep1) or replicate 2 (rep2), “supported by PRICE Calu3” and “supported by ORF-RATER Calu3” whether the ORF was predicted by PRICE or ORF-RATER from the Calu3 data,, the “start position” and “end position” in SAS-CoV-2 genome, the nature of the “start codon”, “size(aa)”and “sequence”.

**Table S5**. Translation levels of SARS-CoV-2 ORFs.

This table lists all translated SARS-CoV-2 ORFs, canonical and newly identified, and their estimated translation levels based on ribosome profiling. “ORF_ID” and “ORF_name” for each ORF, “type” of ORF including upstream ORFs (uORF), in-frame and out-of-frame internal ORFs (iORF and oof), extended versions of canonical ORF (extension) and canonical ORFs. The genomic region used for calculation of coverage is shown in “Included region”. For in-frame iORFs, the coverage in an upstream region of the main ORF, shown in square brackets, was subtracted from the coverage in the included region to get an approximation of expression. See methods for details. Normalized translation levels in VeroE6 cells and in Calu3 cells are shown for each replicate (“chx_1_tpm” and “chx_2_tpm”) and as average value (“chx_mean_tpm”). Alternative calculation of translation levels using ORF-RATER that utilize regression is also presented (“ORF-RATER_chx_1”,” ORF-RATER_chx_2” and “ORF-RATER_chx_mean”).

**Table S6**. Multiple sequence alignment for canonical and novel SARS-CoV-2 ORFs in Sarbecoviruses.

This table includes links for annotated multiple sequence alignment (MSA) views of all SARS-CoV-2, canonical and novel, described in this study, using the CodAlignView tool ^37^. The MSA includes SARS-CoV-2, SARS-CoV and 42 bat coronavirus genomes ^30^.

## Acknowledgements

We thank Stern-Ginossar lab members, Igor Ulitsky and Schraga Schwartz for providing valuable feedback, to Miri Shnayder, Igor Ulitsky and Noa Gil for technical assistance. We thank Emanuel Wyler for the Calu3 cells. We thank Inbar Cohen-Gihon for sharing sequencing and bioinformatics data. This study was supported by the Ben B. and Joyce E. Eisenberg Foundation. Work in the Stern-Ginossar lab is supported by a European Research Council starting grant (StG-2014-638142) and by the Israel Science Foundation (ISF) grant no. 1526/18. S.W-G. is the recipient of the Human Frontier Science Program fellowship (LT-000396/2018), EMBO non-stipendiary Long-Term Fellowship (ALTF 883-2017), the Gruss-Lipper Postdoctoral Fellowship, the Zuckerman STEM Leadership Program Fellowship and the Rothschild Postdoctoral Fellowship. N.S-G is an incumbent of the Skirball Career Development Chair in New Scientists and is a member of the European Molecular Biology Organization (EMBO) Young Investigator Program. The authors declare no competing interests.

## Author contributions

Y.F., O.M., N.P and N.S-G. conceptualization. O.M. experiments. Y.F., A.N. and S.W-G. data analysis. Y.Y-R., H.T., H.A., S.M., S.W., I.C-G, D.S, O.I, A. B-D, T.I. and N.P. work with SARS-CoV-2. D.M mined published proteomic data, Y.F., O.M., A.N., M.S. and N.S-G. interpreted data. M.S. and N.S.-G. wrote the manuscript with contribution from all other authors.

## Material and methods

### Cells and viruses

Vero C1008 (Vero E6) (ATCC CRL-1586™) were cultured in T-75 flasks with DMEM supplemented with 10% fetal bovine serum (FBS), MEM non-essential amino acids, 2mM L-Glutamine, 100Units/ml Penicillin, 0.1mg/ml streptomycin, 12.5Units/ml Nystatin (Biological Industries, Israel). Calu3 cells were cultured in 10cm plates with DMEM supplemented with 10% fetal bovine serum (FBS), MEM non-essential amino acids, 2mM L-Glutamine, 100Units/ml Penicillin, 1% non-essential amino acid and 1% Na-pyrovate. Monolayers were washed once with DMEM (for VeroE6) or RPMI (for Calu3) without FBS and infected with SARS-CoV-2 virus, at a multiplicity of infection (MOI) of 0.2, For calu3 infection 20 ug per ml TPCK trypsin (Thermo scientific) were added. After 1hr infection cells were cultured in their respective medium supplemented with 2% fetal bovine serum, and MEM non-essential amino acids, L glutamine and penicillin-streptomycin-Nystatin at 37°C, 5% CO2. SARS-CoV-2 (GISAID Acc. No. EPI_ISL_406862), was kindly provided by Bundeswehr Institute of Microbiology, Munich, Germany. It was propagated (4 passages) and tittered on Vero E6 cells and then sequenced (details below) before in was used. SARS-CoV-2 BavPat1/2020 Ref-SKU: 026V-03883 was kindly provided by Prof. C. Drosten, Charité – Universitätsmedizin Berlin, Germany. It was propagated (5 passages), tittered on Vero E6 and then sequenced before it has been used in experiments. Infected cells were harvested at the indicated times as described below. Handling and working with SARS-CoV-2 virus was conducted in a BSL3 facility in accordance with the biosafety guidelines of the Israel Institute for Biological Research. The Institutional Biosafety Committee of Weizmann Institute approved the protocol used in these studies.

### Preparation of ribosome profiling and RNA sequencing samples

For RNA-seq, cells were washed with PBS and then harvested with Tri-Reagent (Sigma-Aldrich), total RNA was extracted, and poly-A selection was performed using Dynabeads mRNA DIRECT Purification Kit (Invitrogen) mRNA sample was subjected to DNAseI treatment and 3’ dephosphorylation using FastAP Thermosensitive Alkaline Phosphatase (Thermo Scientific) and T4 PNK (NEB) followed by 3’ adaptor ligation using T4 ligase (NEB). The ligated products used for reverse transcription with SSIII (Invitrogen) for first strand cDNA synthesis. The cDNA products were 3’ ligated with a second adaptor using T4 ligase and amplified for 8 cycles in a PCR for final library products of 200-300bp. For Ribo-seq libraries, cells were treated with either 50μM lactimidomycin (LTM) for 30 minutes or 2μg/mL Harringtonine (Harr) for 5 minutes, for translation initiation libraries (LTM and Harr libraries correspondingly), or left untreated for the translation elongation libraries (cycloheximide [CHX] library). All three samples were subsequently treated with 100μg/mL CHX for 1 minute. Cells were then placed on ice, washed twice with PBS containing 100μg/mL CHX, scraped from the T-75 flasks (Vero cells) or 10cm plates (Calu3 cells), pelleted and lysed with lysis buffer (1% triton in 20mM Tris 7.5, 150mM NaCl, 5mM MgCl_2_, 1mM dithiothreitol supplemented with 10 U/ml Turbo DNase and 100μg/ml cycloheximide). After lysis samples stood on ice for 2h and subsequent Ribo-seq library generation was performed as previously described ^5^. Briefly, cell lysate was treated with RNAseI for 45min at room temperature followed by SUPERse-In quenching. Sample was loaded on sucrose solution (34% sucrose, 20mM Tris 7.5, 150mM NaCl, 5mM MgCl2, 1mM dithiothreitol and 100μg/ml cycloheximide) and spun for 1h at 100K RPM using TLA-110 rotor (Beckman) at 4c. Pellet was harvested using TRI reagent and the RNA was collected using chloroform phase separation. For size selection, 15uG of total RNA was loaded into 15% TBE-UREA gel for 65min, and 28-34 footprints were excised using 28 and 34 flanking RNA oligos, followed by RNA extraction and ribo-seq protocol ^5^

### Virus genomic sequencing

RNA from viruses (culture supernatant after removal of cell debris) was extracted using viral RNA kit (Qiagen). The SMARTer Pico RNA V2 Kit (Clontech) was used for library preparation. Genome sequencing was conducted on the Illumina Miseq platform, in a single read mode 60bp for BetaCoV/Germany/BavPat1/2020 EPI_ISL_406862 and in a paired-end mode 150bp x2 for BavPat1/2020 Ref-SKU: 026V-03883 producing 2,239,263 and 4,332,551 reads correspondingly. Reads were aligned to the viral genome using STAR 2.5.3a aligner. Even coverage along the genome was assessed and the relative abundance junctions (that may reflect genomic deletion) were calculated. For EPI_ISL_406862 passage 4 (that was used for Vero cells infection) the junctions that were found in more than 1% of genomes are listed in Table S2. For BavPat1/2020 Ref-SKU: 026V-03883 passage 5 (that was used to for Calu3 infection) no junctions in abundance of more than 1% of the genomes were detected. All genomic sequencing data was deposited.

### Sequence alignment, normalization and metagene analysis

Sequencing reads were aligned as previously described ^38^. Briefly, linker (CTGTAGGCACCATCAAT) and poly-A sequences were removed and the remaining reads from were aligned to the Chlorocebus sabaeus genome (ENSEMBL release 99) and to the SARS-Cov-2 genomes [Genebank NC_045512.2 with 3 changes to match the used strain (BetaCoV/Germany/BavPat1/2020 EPI_ISL_406862), 241:C->T, 3037:C->T, 23403:A->G]. (infection of Vero cells) or to the Hg19 and NC_045512.2 with the same sequence changes (infection of Calu3). Alignment was performed using Bowtie v1.1.2 ^39^ with maximum two mismatches per read. Reads that were not aligned to the genome were aligned to the transcriptome of Chlorocebus sabaeus (ENSEMBL) and to SARS-CoV-2 junctions that were recently annotated ^18^. The aligned position on the genome was determined as the 5’ position of RNA-seq reads, and for Ribo-seq reads the p-site of the ribosome was calculated according to reads length using the off-set from the 5’ end of the reads that was calculated from canonical cellular ORFs. The offsets used are +12 for reads that were 28-29 bp and +13 for reads that were 30-33 bp. Reads that were in different length were discarded. In all figures presenting ribosome densities data all footprint lengths (28-33bp) are presented.

Novel junctions were mapped using STAR 2.5.3a aligner ^40^, with running flags as suggested at Kim et. al., to overcome filtering of non-canonical junctions. Reads aligned to multiple locations were discarded. Junctions with 5’ break sites mapped to genomic location 55-85 were assigned as leader-dependent junctions. Matching of leader junctions to ORFs, and categorization of junctions as canonical or non-canonical, was adapted from Kim et. al. ^18^ Supplementary table 3, or was assigned manually for strong novel junctions that appear only in our data.

For the metagene analysis only genes with more than 50 reads were used. For each gene normalization was done to its maximum signal and each position was normalized to the number of genes contributing to the position. In the virus 24hr samples, normalization for each gene was done to its maximum signal within the presented region.

For comparing transcript expression level, mRNA and footprint counts from bowtie alignment were normalized to units of RPKM in order to normalize for gene length and for sequencing depth, based on the total number reads for both the host and the virus. The deconvolution of RPKM for RNAs was done by subtracting the RPKM of a gene from the RPKM of the gene located just upstream of it in the genome. The junction counts were based on STAR alignment number of uniquely mapped reads crossing the junction.

The estimation of the viral footprint densities from the 24 hpi samples was performed by calculating the ratio of the RPKM of ORF1a to the total number of leader canonical junctions at 5hpi. This ratio was used as a factor to calculate a proxy for the “true’ viral footprint densities from the number of footprints that were mapped to leader canonical junctions at 24hpi.

To quantify the translation levels of novel viral ORFs at 5hpi and 7hpi, many of which are overlapping, three types of calculations were used based on ORF type. For ORFs that have a unique region, with no overlap to any other ORF, bowtie aligned read density was calculated in that region. For out-of-frame internal ORFs, the read density of the internal ORF region was calculated by estimating the expected 3-bp periodicity distribution of footprints based on non-overlapping translated regions in the main ORF. Using linear regression, we calculated the relative contribution of the frames of the main and of the internal ORF to the reads covering the region of the internal ORF. The relative contribution of the internal ORF was then multiplied by the read density in that region to obtain the estimated translation level of the internal out-of-frame ORF. For in-frame internal ORFs the read density of the main overlapping ORF is calculated from a non-overlapping region and then subtracted from the read density in the overlapping internal ORF region to get an estimate of translation levels of the internal ORF. In cases where the unique region used to calculate read density contained the start-codon of the ORF, the first 20% of the codons in the region were excluded from the calculation to avoid bias from initiation peaks, unless the region was very short and trimming it would harm the ability to estimate coverage (ORF 8 and extended ORF M). The exact regions that were used for calculation can be found in Table S5. Finally, read density was normalized to the length of the region used for calculation and to the sum of length normalized reads in each sample to get TPM values. P-values for the relative contribution levels of out-of-frame ORFs were calculated from both replicates using a mixed-effects linear model using the 3-base periodicity distribution as the fixed effect and the replicates as random effect. In parallel, ORF-RATER was used to quantify the translation levels of the viral ORFs (using regression), giving largely similar values (Spearmen’s R = 0.92 and R = 0.87 in VeroE6 and Calu3, respectively).

### Prediction of translation initiation sites and transmembrane domains

Translation initiation sites were predicted using PRICE ^24^ and ORF-RATER ^25^. To estimate the codons generating the sequencing reads with maximum likelihood, PRICE requires a predefined set of annotated coding sequences from the same experiment. Thus, it does not perform well on reference sequences with a small number of annotated ORFs such as SARS-CoV-2. Since our experiment generated ribosome footprints from both SARS-CoV-2 and host mRNAs, which were exposed to the exact same conditions in the protocol, we used annotated CDSs from the host cells to evaluate the parameters of the experiment. For libraries of infected Vero cells sequencing reads were aligned using Bowtie to a fasta file containing chromosome 20 of Chlorocebus sabaeus (1240 annotated start codons, downloaded from ensembl: ftp://ftp.ensembl.org/pub/release99/fasta/chlorocebus_sabaeus/dna/) and the genomic sequence of SARS-CoV-2 (Refseq NC_045512.2). A gtf file with the annotations of Chlorocebus sabaeus and SARS-CoV-2 genomes was constructed and provided as the annotations file when running PRICE. For technical reasons, the annotation of the first coding sequence (CDS) of the two CDSs in the “ORF1ab” gene was deleted since having two CDSs encoded from a single gene was not permitted by PRICE. For libraries of infected Calu3 cells sequencing reads were mapped to a fasta file containing chromosome 1 of hg19 (2843 annotated start codons) and the genomic sequence of SARS-CoV-2 (Refseq NC_045512.2). A gtf file with the annotations of hg19 and SARS-CoV-2 genomes was constructed and provided as the annotations file when running PRICE. For the data that was generated from infected Vero cells at 5hpi training and ORF prediction by PRICE were done once using the CHX data from both replicates, and again using all Ribo-seq libraries from both replicates, and the resulting predictions were combined. To test reproducibility, the same predictions were performed on each replicate separately. For the data that was generated from infected Calu3 cells at 7hpi training and ORF prediction by PRICE were done using all Ribo-seq libraries from both replicates The predictions were further filtered to include only ORFs with at least 100 reads at the initiation site in the LTM samples of at least one replicate. ORFs were then defined by extending each initiating codon to the next in-frame stop codon.

ORF-RATER was used with the default values besides allowing all start codons with at most one mismatch to ATG. For each cell type, two runs of ORF-RATER were used. One in which ORF-RATER was trained on cellular annotations (chr 20 for the Vero cells, and chr 1 for the Calu cells) and SARS-CoV-2 canonical ORFs (similar to the procedure that was used for running PRICE). In the second run only SARS-CoV-2 canonical ORFs were used for training. In both cases ORF1b and ORF10 were omitted from the training set. BAM files from STAR alignment were used as input. The CHX data from both replicates was used in the first prune step to omit low coverage ORFs. The calculations of the P-site offsets, and the regression were performed for each Ribo-Seq library separately. The final score was calculated based on all three types of libraries. Score of 0.5 was used as cut-off for the final predictions these were further manually curated. Additional ORFs that were not recognized by the trained models (likely due to differences in the features of viral genome compared to cellular genomes) but presented reproducible translation profile in the two cell lines were added manually to the final ORF list (Table S4). ORFs were manually identified as such if they had reproducible initiation peaks in the CHX libraries that were enhanced in the LTM and Harr libraries, and exhibited increased CHX signal in the correct reading-frame along the coding region.

### Mapping reads to CUG initiation upstream the TRS-leader

Reads from ribosome profiling libraries were aligned using bowtie to a single reference that contained the transcripts and the genome allowing no mismatches or gaps. Reads with p-site mapped to position 59 of the viral genome were collected and divided to four groups according to the nucleotide in position +17 of the read (position 76 of the genome). The first group contains reads that are short (28 nucleotides) and do not have any nucleotide at position +17. The other three groups, referred to as T, A and G, correspond to combinations of genomic and subgenomic RNAs based on their sequence, as shown in figure S14. Group T is attributed to the genome or to ORF E and ORF M subgenomic RNAs, group A to the subgenomic RNAs of ORF S, ORF7a, ORF8 and ORF N, and group G to the sub-genomic RNA of ORF 6. Reads mapped uniquely to the subgenomic RNA of ORF3a were excluded from calculation, and the number of reads in each group was summed. Group T, containing genomic reads, was further divided based on the nucleotide at position +18, where reads with A at that position can originate from the subgenomic RNA of ORF M and reads with T at that position can originate from the genome or from the subgenomic RNA of ORF E. Final division of the genomic group was done based on position +19 where T corresponds to genomic reads and A corresponds to ORF E subgenomic reads. RNA values as calculated from junction densities (described above) were summed for the subgenomic and genomic RNAs in each group. The analysis was performed for each ribosome profiling library separately.

### Mining of proteomics data

Data downloaded from Bojkova et al.14 was searched using Byonic search engine using 10ppm tolerance for MS1 and 20ppm tolerance for MS2, against the concatenated database containing our 26 novels ORFs as well as the human proteome DB (SwissProt Nov2019), and the SARS-CoV-2 proteome. Modifications allowed were fixed carbamidomethylation on C, fixed TMT6 on K and peptide N-terminus, variable K8 and R10 SILAC labeling, variable M oxidation and Variable NQ deamidation. Data downloaded from Davidson et al.15 was searched using Byonic search engine using 10ppm tolerance for MS1 and 0.6Da tolerance for MS2, against the concatenated database containing our 26 novel ORFs as well as the human proteome DB (SwissProt Nov2019), and the SARS-CoV-2 proteome. Modifications allowed were fixed carbamidomethylation on C, variable N-terminal protein acetylation, M oxidation and NQ deamidation.

### Immunofluorescence

Cells were plated on ibidi slides, fixed in 3% paraformaldehyde for 20 minutes, permeabilized with 0.5% Triton X-100 in PBS for 2 minutes, and then blocked with 2% FBS in PBS for 30 minutes. Immunostaining was performed with rabbit anti-SARS-CoV-2 serum. Cells were washed and labeled with anti-rabbit FITC antibody and with DAPI (4=,6-diamidino-2-phenylindole). Imaging was performed on a Zeiss AxioObserver Z1 wide-field microscope using a X40 objective and Axiocam 506 mono camera.

### Data availability

All next-generation sequencing data files were deposited in Gene Expression Omnibus under accession number GSE149973.

All the RNA-seq and ribosome profiling data generated in this study can be accessed through a UCSC browser session: http://genome.ucsc.edu/s/aharonn/CoV2%2DTranslation

## References

1. Zhu, N. et al. A novel coronavirus from patients with pneumonia in China, 2019. N. Engl. J. Med. 382, 727–733 (2020).

2. Zhou, P. et al. A pneumonia outbreak associated with a new coronavirus of probable bat origin. Nature 579, 270–273 (2020).

3. Ingolia, N. T., Lareau, L. F. & Weissman, J. S. Ribosome profiling of mouse embryonic stem cells reveals the complexity and dynamics of mammalian proteomes. Cell 147, 789–802 (2011).

4. Lee, S. et al. Global mapping of translation initiation sites in mammalian cells at single-nucleotide resolution. Proc. Natl. Acad. Sci. U. S. A. 109, E2424–E2432 (2012).

5. Finkel, Y. et al. Comprehensive annotations of human herpesvirus 6A and 6B genomes reveal novel and conserved genomic features. Elife 9, e50960 (2020).

6. Stern-Ginossar, N. et al. Decoding human cytomegalovirus. Science (80-.). 338, 1088–1093 (2012).

7. Irigoyen, N. et al. High-Resolution Analysis of Coronavirus Gene Expression by RNA Sequencing and Ribosome Profiling. PLoS Pathog. 12, 1005473 (2016).

8. Dinan, A. M. et al. Comparative Analysis of Gene Expression in Virulent and Attenuated Strains of Infectious Bronchitis Virus at Subcodon Resolution. J. Virol. 93, 714–733 (2019).

9. Snijder, E. J., Decroly, E. & Ziebuhr, J. The Nonstructural Proteins Directing Coronavirus RNA Synthesis and Processing. in Advances in Virus Research 96, 59–126 (Academic Press Inc., 2016).

10. Sola, I., Almazán, F., Zúñiga, S. & Enjuanes, L. Continuous and Discontinuous RNA Synthesis in Coronaviruses. Annu. Rev. Virol. 2, 265–288 (2015).

11. Stadler, K. et al. SARS — beginning to understand a new virus. Nat. Rev. Microbiol. 1, 209–218 (2003).

12. Lai, M. M. & Stohlman, S. A. Comparative analysis of RNA genomes of mouse hepatitis viruses. J. Virol. 38, 661–670 (1981).

13. Yogo, Y., Hirano, N., Hino, S., Shibuta, H. & Matumoto, M. Polyadenylate in the virion RNA of mouse hepatitis virus. Journal of Biochemistry 82, (1977).

14. Bojkova, D. et al. Proteomics of SARS-CoV-2-infected host cells reveals therapy targets. (2020). doi:10.1038/s41586-020-2332-7

15. Davidson, A. D. et al. Characterisation of the transcriptome and proteome of SARS-CoV-2 using direct RNA sequencing and tandem mass spectrometry reveals evidence for a cell passage induced in-frame deletion in the spike glycoprotein that removes the furin-like cleavage site. bioRxiv 2020.03.22.002204 (2020). doi:10.1101/2020.03.22.002204

16. Ingolia, N. T. et al. Ribosome Profiling Reveals Pervasive Translation Outside of Annotated Protein-Coding Genes. Cell Rep. 8, 1365–1379 (2014).

17. Plant, E. P., Rakauskaite, R., Taylor, D. R. & Dinman, J. D. Achieving a golden mean: mechanisms by which coronaviruses ensure synthesis of the correct stoichiometric ratios of viral proteins. J. Virol. 84, 4330–40 (2010).

18. Kim, D. et al. The architecture of SARS-CoV-2 transcriptome. Cell S0092-8674, 30406–2 (2020).

19. Schaecher, S. R., Mackenzie, J. M. & Pekosz, A. The ORF7b protein of severe acute respiratory syndrome coronavirus (SARS-CoV) is expressed in virus-infected cells and incorporated into SARS-CoV particles. J. Virol. 81, 718–31 (2007).

20. Liu, Z. et al. Identification of a common deletion in the spike protein of SARS-CoV-2. bioRxiv 2020.03.31.015941 (2020). doi:10.1101/2020.03.31.015941

21. Ogando, N. S. et al. SARS-coronavirus-2 replication in Vero E6 cells: replication kinetics, rapid adaptation and cytopathology. bioRxiv 2020.04.20.049924 (2020). doi:10.1101/2020.04.20.049924

22. Taiaroa, G. et al. Direct RNA sequencing and early evolution of SARS-CoV-2. bioRxiv 2020.03.05.976167 (2020). doi:10.1101/2020.03.05.976167

23. Wu, A. et al. Genome Composition and Divergence of the Novel Coronavirus (2019-nCoV) Originating in China. Cell Host Microbe 27, 325–328 (2020).

24. Erhard, F. et al. Improved Ribo-seq enables identification of cryptic translation events. Nat. Methods 15, 363–366 (2018).

25. Fields, A. P. et al. A Regression-Based Analysis of Ribosome-Profiling Data Reveals a Conserved Complexity to Mammalian Translation. Mol. Cell 60, 816–827 (2015).

26. Bouhaddou, M. et al. The Global Phosphorylation Landscape of SARS-CoV-2 Infection. Cell (2020). doi:10.1016/j.cell.2020.06.034

27. Stukalov, A. et al. Multi-level proteomics reveals host-perturbation strategies of SARS-CoV-2 and SARS-CoV. bioRxiv 2020.06.17.156455 (2020). doi:10.1101/2020.06.17.156455

28. Cagliani, R., Forni, D., Clerici, M. & Sironi, M. Coding potential and sequence conservation of SARS-CoV-2 and related animal viruses. Infect. Genet. Evol. 83, (2020).

29. Firth, A. E. A putative new SARS-CoV protein, 3a*, encoded in an ORF overlapping ORF3a. bioRxiv 2020.05.12.088088 (2020). doi:10.1101/2020.05.12.088088

30. Jungreis, I., Sealfon, R. & Kellis, M. Sarbecovirus comparative genomics elucidates gene content of SARS-CoV-2 and functional impact of COVID-19 pandemic mutations. bioRxiv 2020.06.02.130955 (2020). doi:10.1101/2020.06.02.130955

31. Gordon, D. E. et al. A SARS-CoV-2 protein interaction map reveals targets for drug repurposing. Nature 583, 459–468 (2020).

32. Hachim, A. et al. Beyond the Spike: identification of viral targets of the antibody response to SARS-CoV-2 in COVID-19 patients. medRxiv 2020.04.30.20085670 (2020). doi:10.1101/2020.04.30.20085670

33. Yewdell, J. W. DRiPs solidify: Progress in understanding endogenous MHC class I antigen processing. Trends in Immunology 32, 548–558 (2011).

34. Abernathy, E. & Glaunsinger, B. Emerging roles for RNA degradation in viral replication and antiviral defense. Virology 479-480, 600–608 (2015).

35. Huang, C. et al. SARS coronavirus nsp1 protein induces template-dependent endonucleolytic cleavage of mRNAs: Viral mRNAs are resistant to nsp1-induced RNA cleavage. PLoS Pathog. 7, e1002433 (2011).

36. Kamitani, W., Huang, C., Narayanan, K., Lokugamage, K. G. & Makino, S. A two-pronged strategy to suppress host protein synthesis by SARS coronavirus Nsp1 protein. Nat. Struct. Mol. Biol. 16, 1134–1140 (2009).

37. Jungreis, I. et al. Evolutionary Dynamics of Abundant Stop Codon Readthrough. Mol. Biol. Evol. 33, 3108–3132 (2016).

38. Tirosh, O. et al. The Transcription and Translation Landscapes during Human Cytomegalovirus Infection Reveal Novel Host-Pathogen Interactions. PLoS Pathog. 11, e1005288 (2015).

39. Langmead, B., Trapnell, C., Pop, M. & Salzberg, S. L. Ultrafast and memory-efficient alignment of short DNA sequences to the human genome. Genome Biol. 10, R25 (2009).

40. Dobin, A. et al. STAR: ultrafast universal RNA-seq aligner. Bioinformatics 29, 15–21 (2013).

