## Supplementary figures for "The coding capacity of SARS-CoV-2"

Figure S1

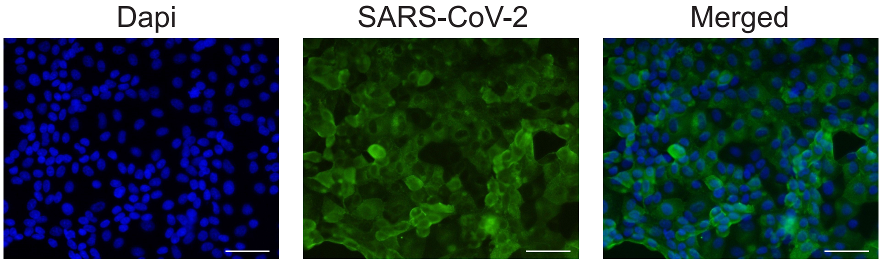


**Figure S1**. 24 hr infection with SARS-CoV-2 of Vero E6 cells.
Vero E6 cells were infected with SARS-CoV-2 at an MOI=0.2 and 24 hpi the cells were fixed and stained with antisera against SARS-CoV-2 (green) and Dapi (blue).
Representative microscopy images are presented. Scale bars are 200μm.

### Figure S2

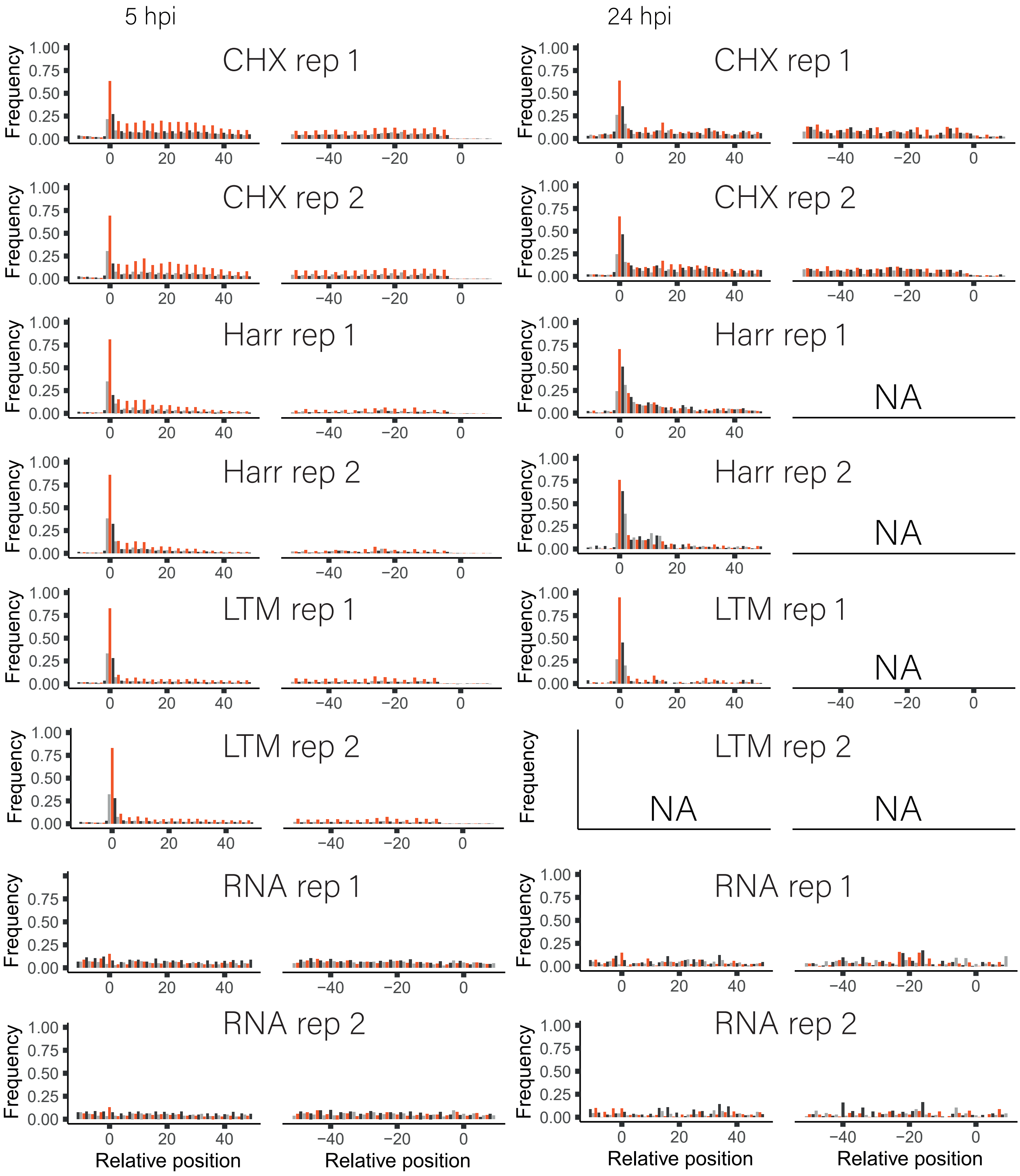


**Figure S2.** Footprint profiles of cellular genes from SARS-CoV-2 infected Vero E6 cells

Metagene analysis of read densities at the 5' and the 3' regions of cellular protein coding genes as measured by the different ribosome profiling approaches and RNA-seq at 5hpi and 24hpi, from two biological replicates. The X axis shows the nucleotide position relative to the start or the stop codons. The ribosome densities are shown with different colors indicating the three frames relative to the main ORF (red, frame 0; black, frame +1; grey, frame +2). NA reflect samples in which we did not obtain enough cellular genes that contain 50 reads at the 5' or 3' regions to generate metagene profile.

Figure S3

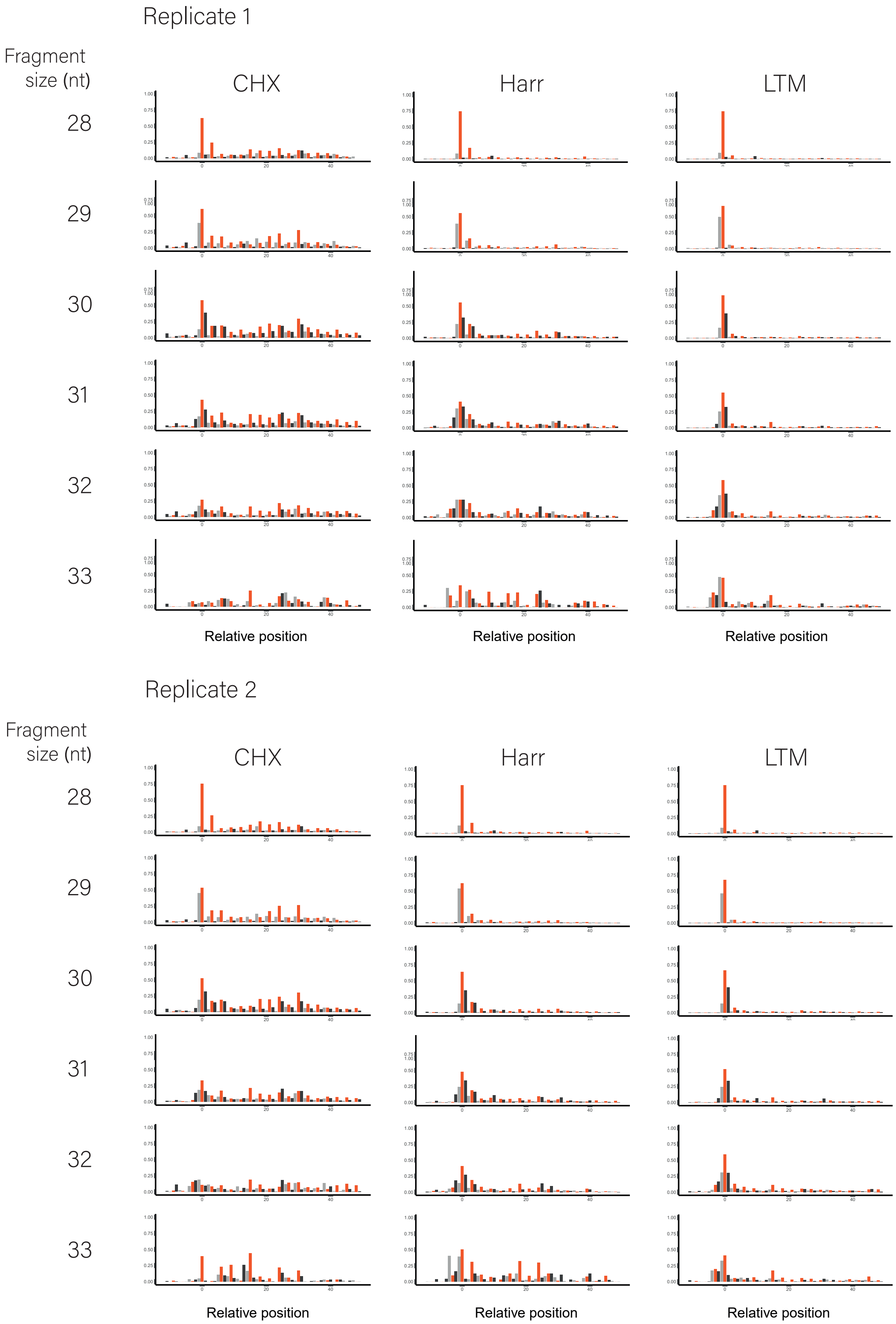


**Figure S3.** Footprint profiles separated based on the protected fragment size

Metagene analysis of read densities at the 5' regions of cellular protein coding genes as measured by the different ribosome profiling approaches at 5hpi, from two biological replicates. The X axis shows the nucleotide position relative to the start or the stop codons. The ribosome densities for each fragment size (28bp-33bp) are shown with different colors indicating the three frames relative to the main ORF (red, frame 0; black, frame +1; grey, frame +2). This analysis indicates that in both replicates 28-29bp footprints show the strongest bias towards the translated frame.

Figure S4

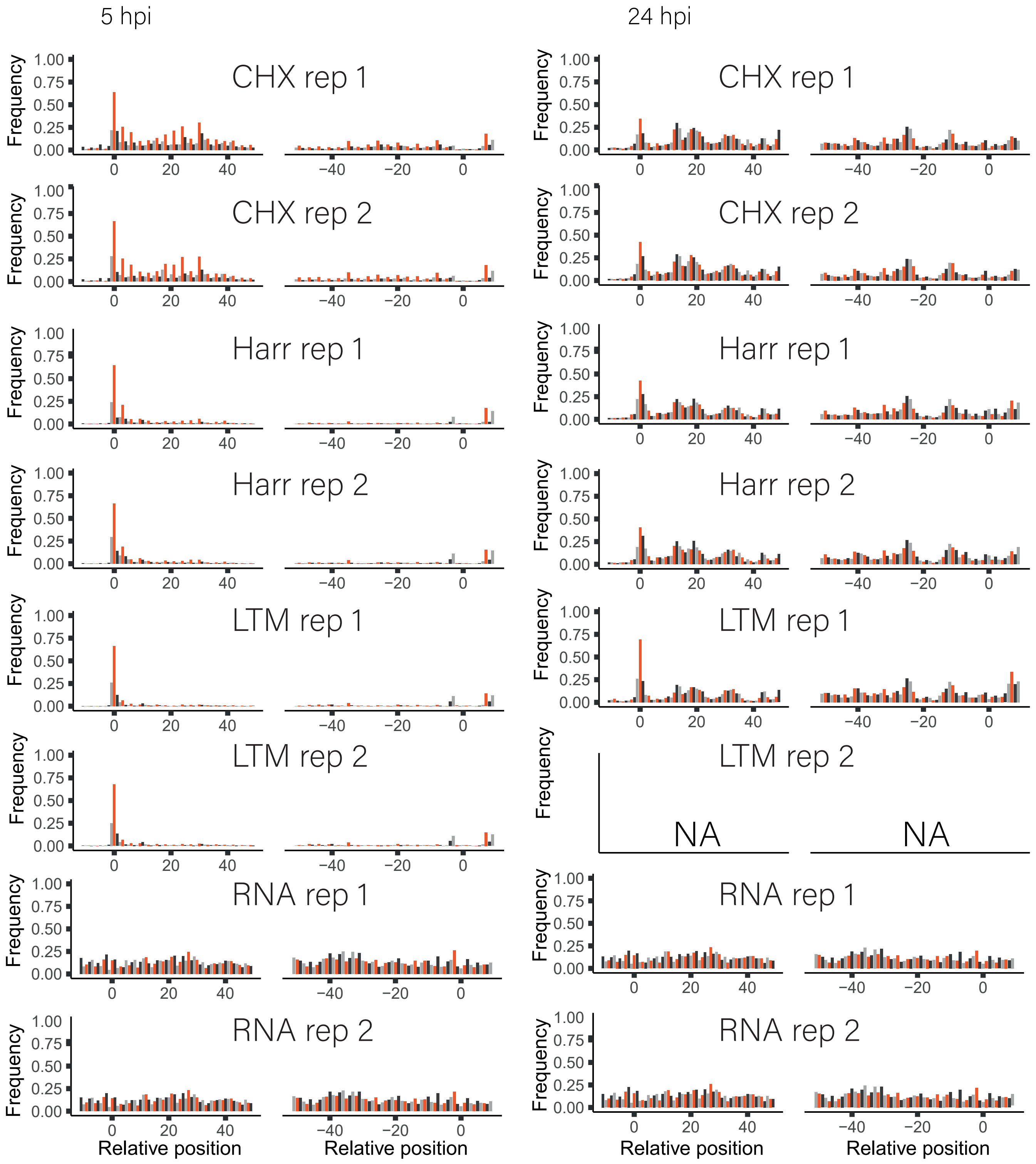


**Figure S4.** Footprint profiles of viral genes from SARS-CoV-2 infected Vero E6 cells

Metagene analysis of read densities at the 5' and the 3' regions of viral protein coding genes as measured by the different ribosome profiling approaches and RNA-seq at 5hpi and 24hpi, from two biological replicates. The X axis shows the nucleotide position relative to the start or the stop codons. The ribosome densities are shown with different colors indicating the three frames relative to the main ORF (red, frame 0; black, frame +1; grey, frame +2). NA reflect samples in which we did not obtain enough reads at the 5' and the 3' regions.

Figure S5

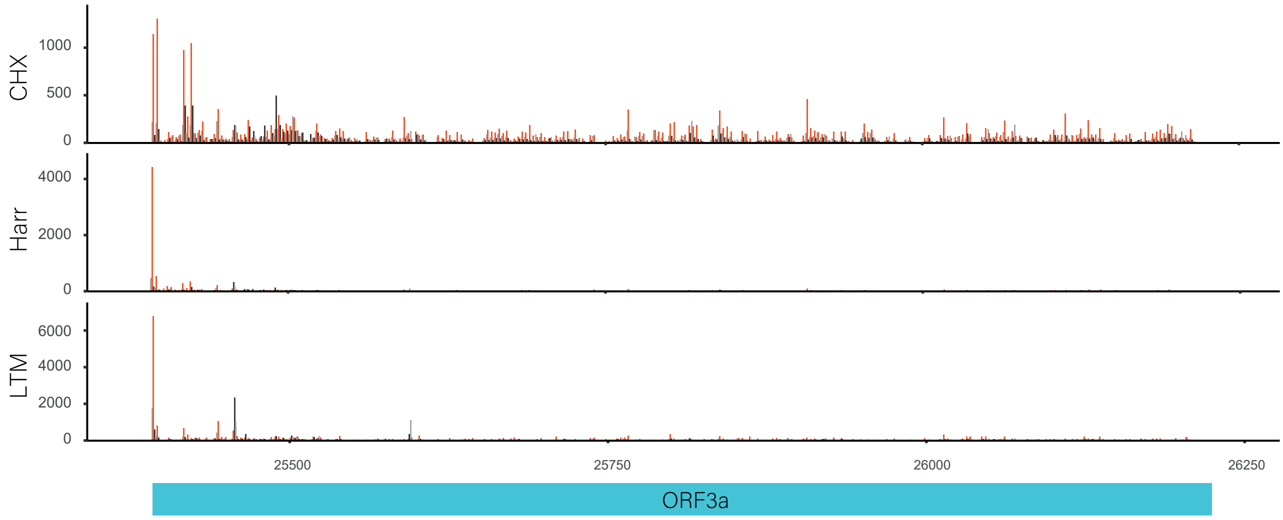


**Figure S5.** Footprint profiles of ORF3a

Read densities over SARS-CoV-2 ORF3a gene as measured by the different ribosome profiling approaches at 5hpi. The X axis shows the nucleotide position on the SARS-CoV-2 genome. The ribosome densities are shown with different colors indicating the three frames relative to the main ORF (red, frame 0; black, frame +1; grey, frame +2).

Figure S6

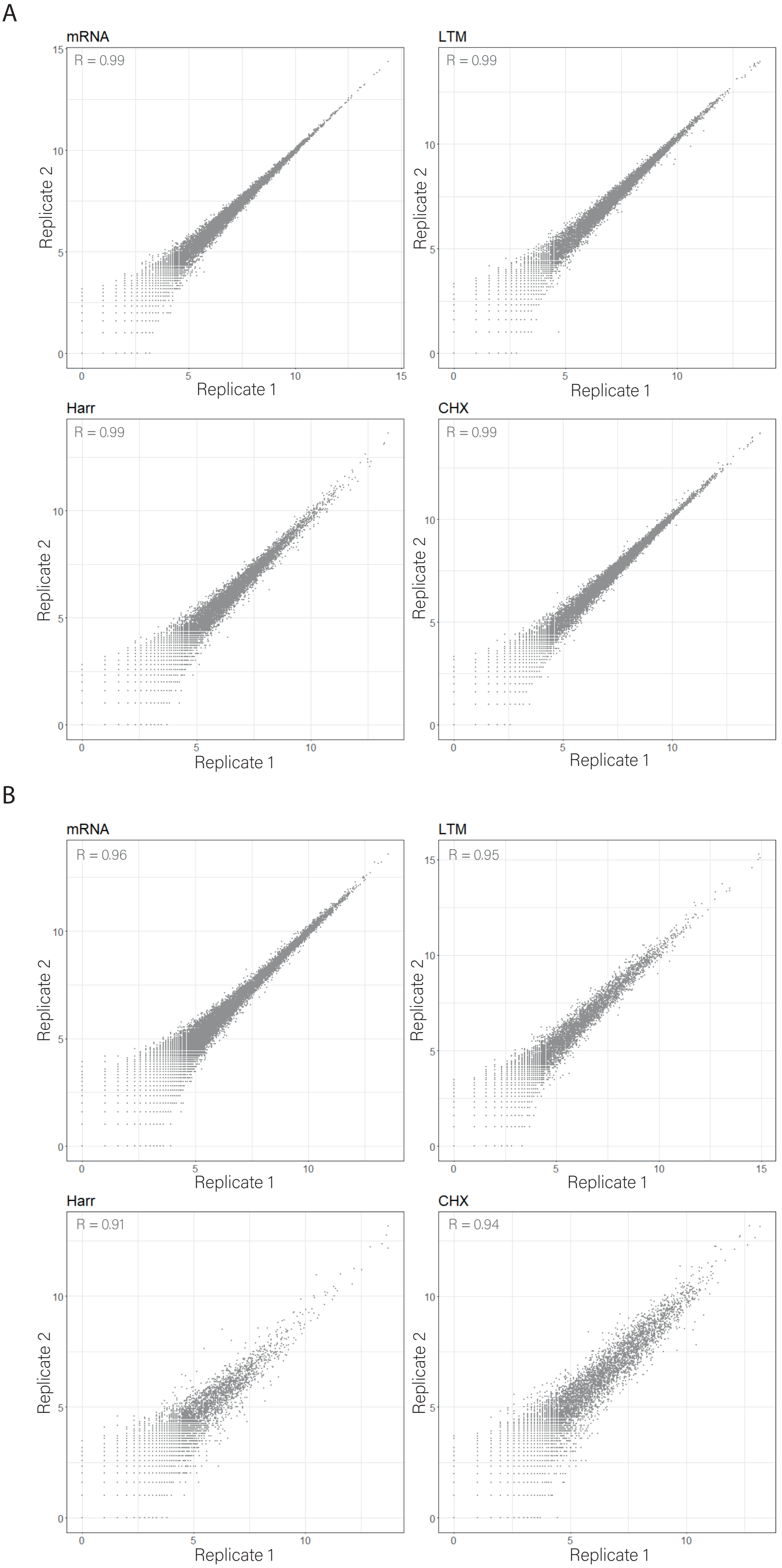


Figure S6. Reproducibility between biological replicates
(A) Scatter plots depicting read densities on cellular gene as measured in two independent biological replicates for mRNA, LTM, HARR and CHX libraries. (B) Scatter plots depicting the number of reads in every position along the SARS-CoV-2 genome in two independent biological replicates, demonstrating reproducibility between our replicates at single codon resolution. Pearson's R of log transformed values is presented.

Figure S7

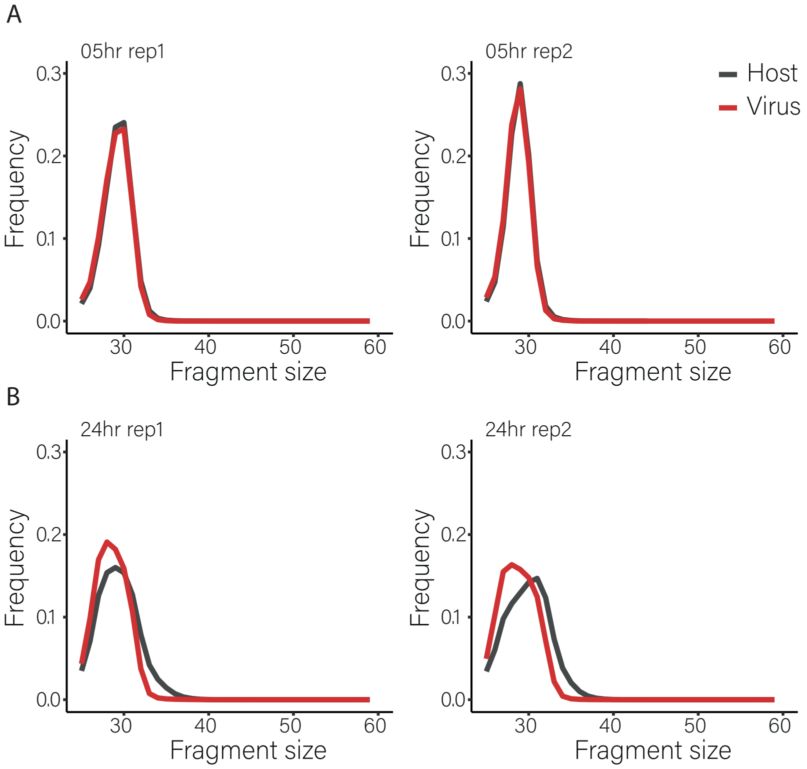


**Figure S7.** Length distribution of ribosome footprint fragments

Length distribution of ribosome footprint fragments from SARS-CoV-2 genes (red) or cellular genes (black) at 5hpi **(A)** and 24hpi **(B)** from both biological replicates.

Figure S8

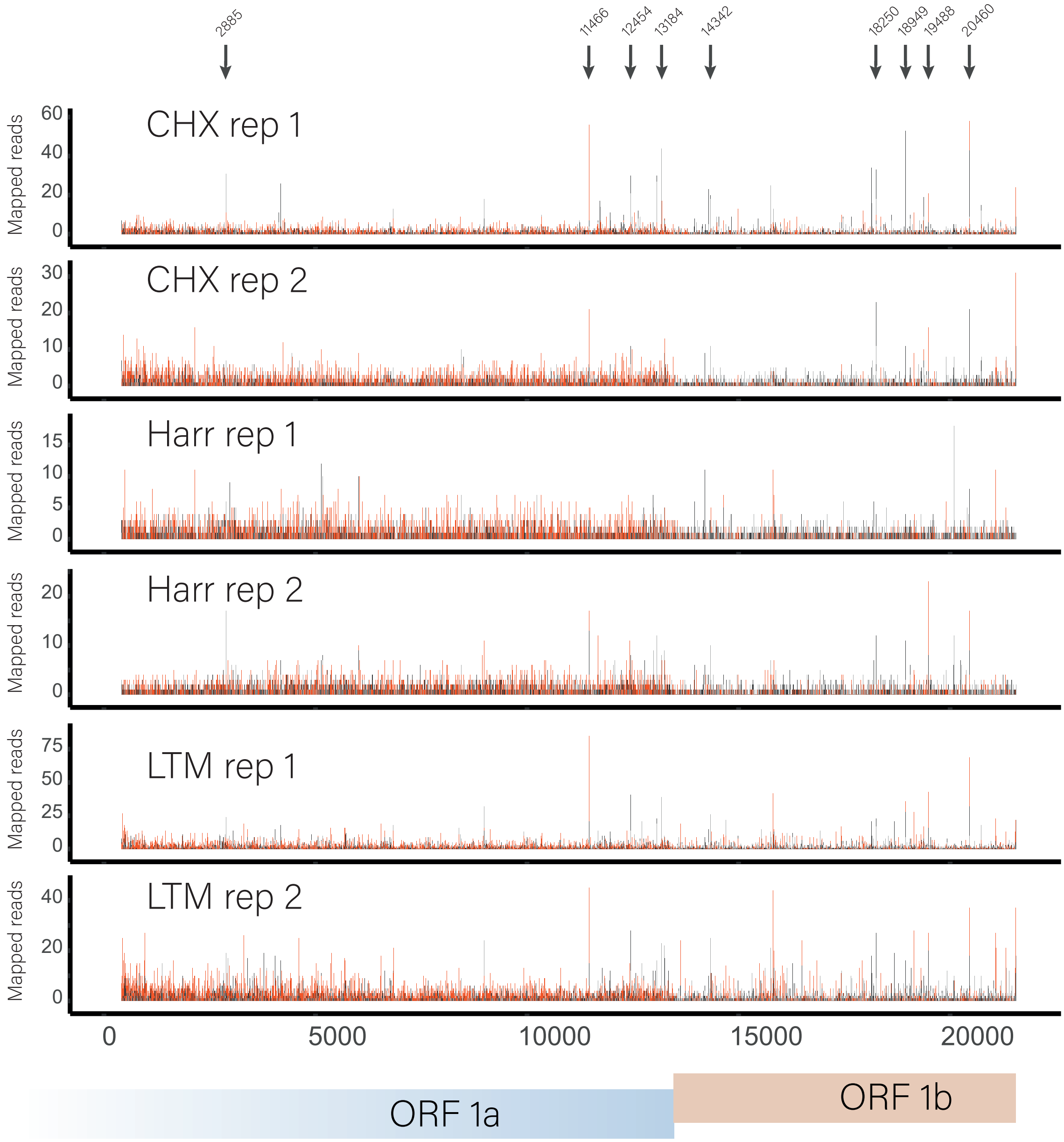


**Figure S8.** Footprint profiles reveal potential ribosome pausing sites within ORF1a and ORF1b

Read densities are presented for ORF1a and ORF1b at 5 hpi from two biological replicates. The ribosome densities are shown with different colors indicating the three frames relative to the translated frame of ORF1a (red, frame 0; black, frame +1; grey, frame +2). Black arrows mark potential ribosome pausing sites and their genomic positions.

Figure S9

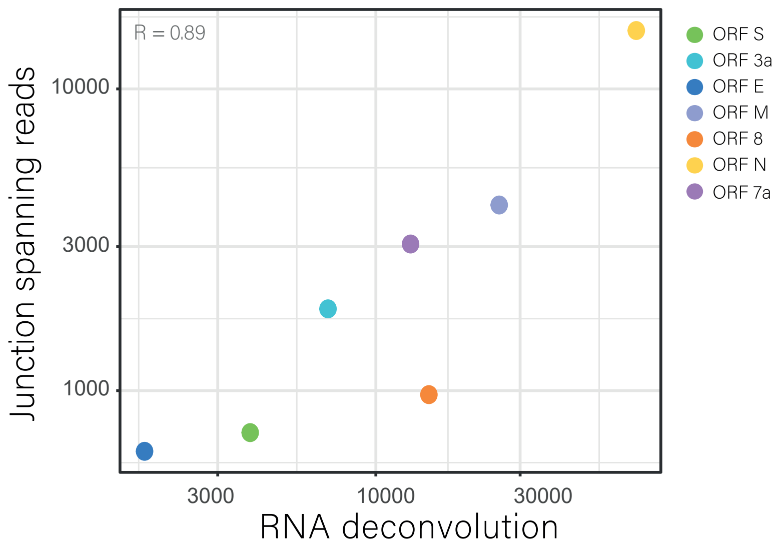


**Figure S9.** Correlation between two approaches for calculating the abundance of subgenomic RNAs

Measurement of subgenomic RNA abundance using deconvolution of RNA densities versus using relative abundance of RNA reads spanning leader-body junctions, for seven canonical viral ORFs. Spearman's R is presented.

Figure S10

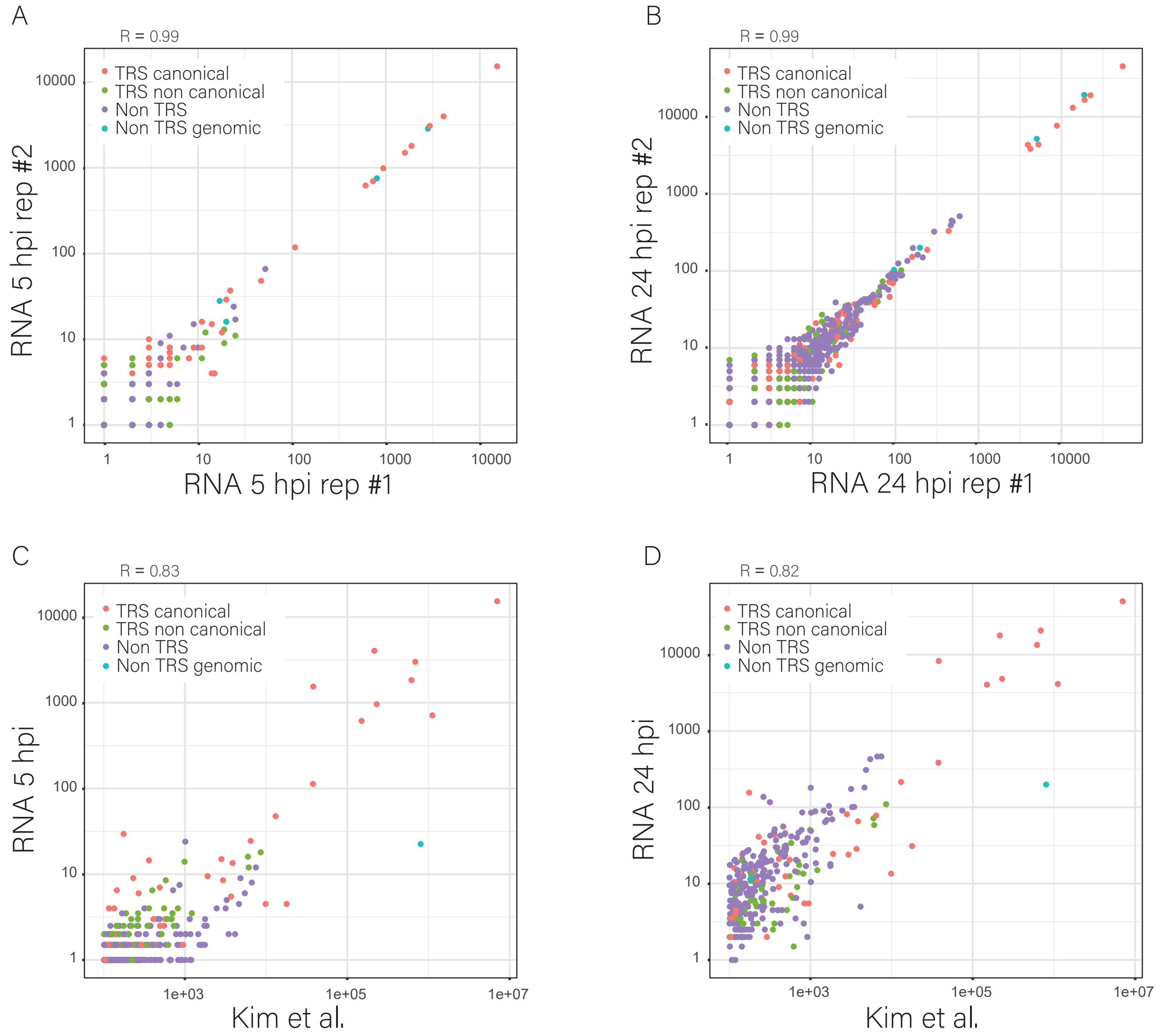


**Figure S10.** Correlation between junction-spanning RNA-seq reads from different samples

Scatter plots presenting the abundance of junction-spanning RNA-seq reads from two biological replicates from **(A)** 5hpi and **(B)** 24hpi. Scatter plots presenting the average abundance of junction-spanning RNA-seq reads from **(C)** 5hpi or **(D)** 24hpi versus data from Kim et al. Viral reads that span canonical leader dependent junctions are marked in red, non-canonical leader dependent junctions are marked in green, non-canonical leader independent junctions are marked in purple and non-canonical leader independent junctions originating from genomic deletions are marked in cyan. Pearson’s R of log transformed values is presented.

Figure S11

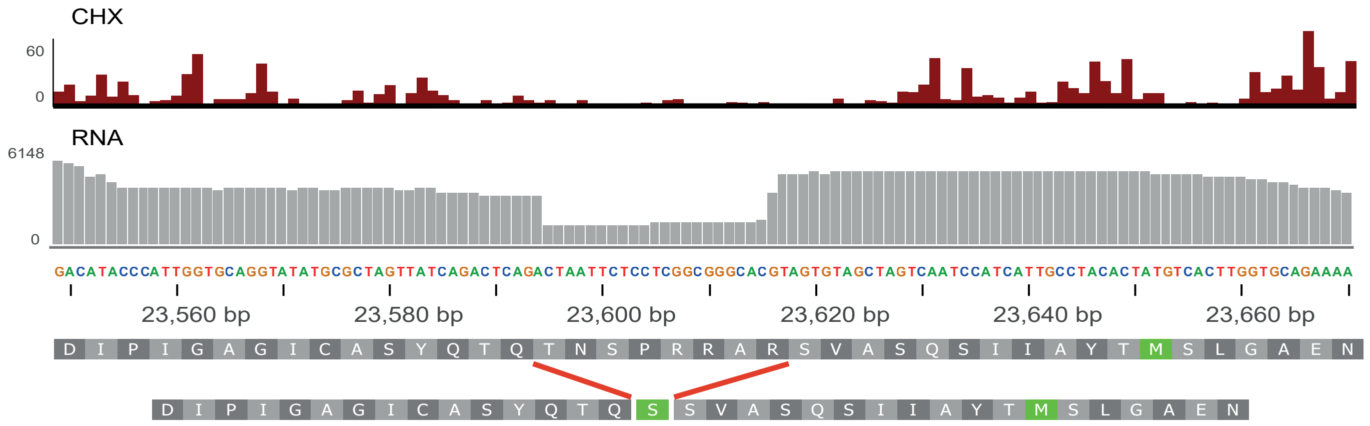


**Figure S11.** An abundant deletion within the Spike protein

Ribosome profiling (CHX) and RNA densities over the 7aa deletion in the S protein. Lower panels present the amino acid sequence in the region and the translation of the WT and of the 7aa deleted version of the S protein.

Figure S12 - page 1

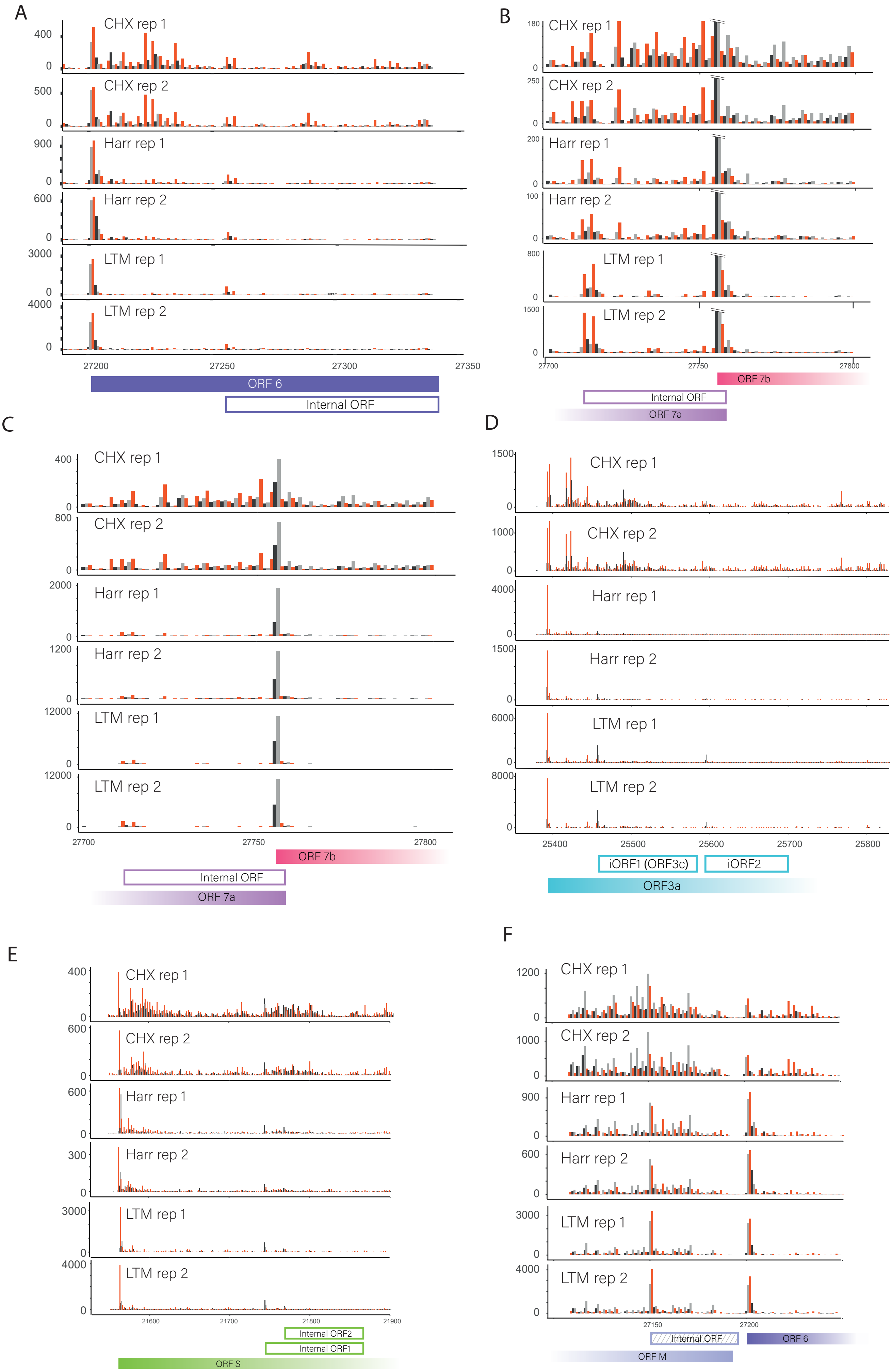

Figure S12 - page 2

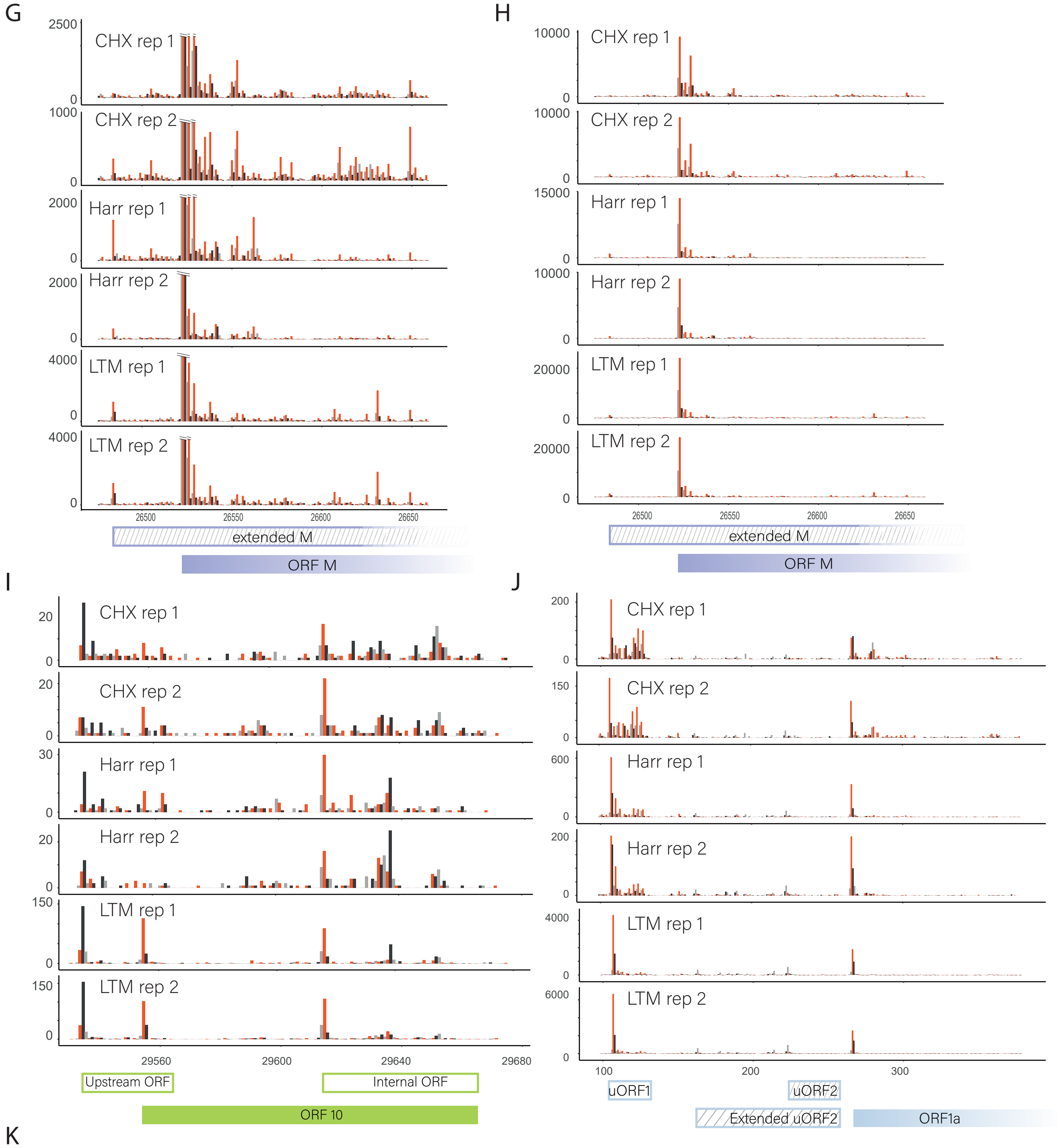


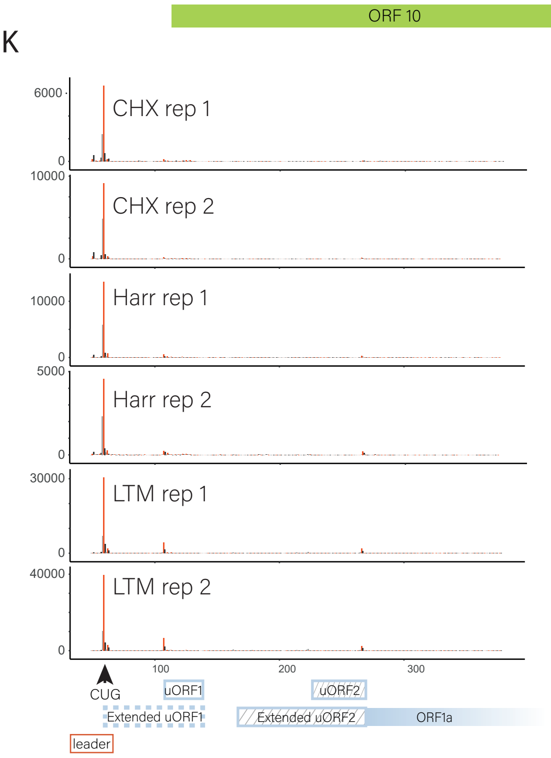


**Figure S12.** Footprint profiles reveal novel viral coding regions (A-K) Read densities over SARS-CoV-2 unannotated translated ORFs as measured by the different ribosome profiling approaches from both replicates at 5hpi. The ribosome densities are shown with different colors indicating the three frames relative to the main ORF in each figure (red, frame 0; black, frame +1; grey, frame +2). Filled and open rectangles indicate the canonical and novel ORFs, respectively. ORFs starting in near cognate start codon are labeled with stripes. (A) In frame internal initiation within ORF6 generating truncated product. (B and C) In frame internal initiation within ORF7a, (B) scale is adapted to present the internal ORF initiation (C) full scale presenting ORF7b initiation in true scale. (D) Out of frame internal initiations within ORF3a. (E) Out of frame internal initiations within ORF-S. (F) Out of frame internal initiation within ORF-M. (G-H) an extended version of ORF-M (G) scale is adapted to present the extended ORF (H) full scale presenting the dominance of the main initiation. (I) uORF that overlaps ORF10 initiation and in frame internal initiation generating truncated ORF10 product (I) two uORFs embedded in ORF1ab 5'UTR (J) non-canonical CUG initiation upstream of the TRS leader

Figure S13

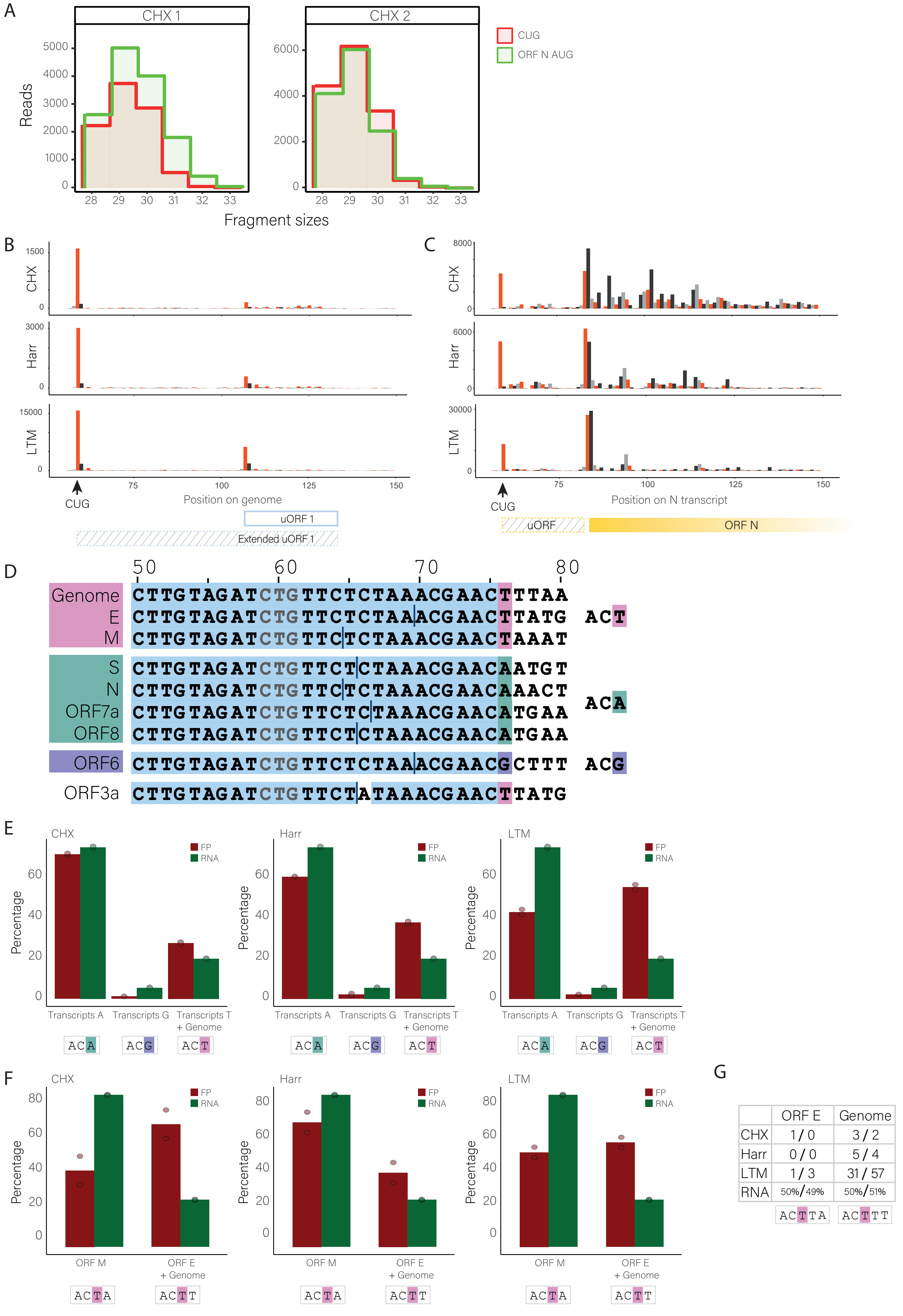
A
CHX 1
CHX 2
CUG
ORF N AUG
Reads
Fragment sizes
B
C
CHX
Harr
LTM
CUG
Position on genome
Position on N transcript
uORF 1
Extended uORF 1
uORF
ORF N
D
Genome
E
M
S
N
ORF7a
ORF8
ORF6
ORF3a
CTTGTAGATCTGTTCTCTAAACGAACTTTAA
CTTGTAGATCTGTTCTCTAAACGAACTTATG
CTTGTAGATCTGTTCTCTAAACGAACTAAAT
CTTGTAGATCTGTTCTCTAAACGAACAATGT
CTTGTAGATCTGTTCTCTAAACGAACAAACT
CTTGTAGATCTGTTCTCTAAACGAACATGAA
CTTGTAGATCTGTTCTCTAAACGAACATGAA
CTTGTAGATCTGTTCTCTAAACGAACGCTTT
CTTGTAGATCTGTTCTATAAACGAACTTATG
ACT
ACA
ACG
E
F
FP
RNA
Percentage
Transcripts A
Transcripts G
Transcripts T + Genome
ORF M
ORF E + Genome
ACTA
ACTT
G
| | ORF E | Genome |
| CHX | 1 / 0 | 3 / 2 |
| Harr | 0 / 0 | 5 / 4 |
| LTM | 1 / 3 | 31 / 57 |
| RNA | 50%/49% | 50%/51% |
ACTTA
ACTTT

**Figure S13.** The CUG initiation is enriched on the genomic RNA

**(A)** Comparison of the ribosome footprint read length distributions of reads that align to the CUG upstream of the TRS-L (red) and reads that align to ORF-N AUG (green). **(B and C)** Read densities over the TRS-L CUG on the genomic RNA **(B)** or ORF-N transcript **(C)** as measured by the different ribosome profiling approaches at 5hpi. The ribosome densities are shown with different colors indicating the three frames relative to the CUG (red, frame 0; black, frame +1; grey, frame +2). Rectangles indicate adjacent ORFs and the striped rectangles indicate the ORFs initiating at the TRS-L CUG. **(D)** The sequences of the genome and the most abundant subgenomic transcripts divided to three groups based on the base in position 76. The CUG position is labeled in gray and the location of the junction is labeled by a vertical line. **(E)** The relative number of footprint reads that their P-site was mapped to the CUG (dark red) for each of the transcript groups (as defined in C) and the relative RNA abundance of these transcript groups (green). Data is presented for CHX, Harr and LTM libraries. **(F)** The relative number of footprint reads that their P-site was mapped to the CUG (dark red) in ORF-M transcript or in ORF-E and the Genome RNA and the relative RNA abundance (green). Only footprint sizes 31-33bp that allow unique alignment were used. Data is presented for CHX, Harr and LTM libraries. **(G)** The number of footprint reads that their P-site was mapped to the CUG in the genome or in ORF-E transcript out of all reads and the relative abundance of these RNAs. Only footprint sizes 32-33bp that allow unique alignment were used. Read numbers is presented for CHX, Harr and LTM libraries and the relative RNA abundance is presented as percentage of total RNA included in the comparison.

Figure S14

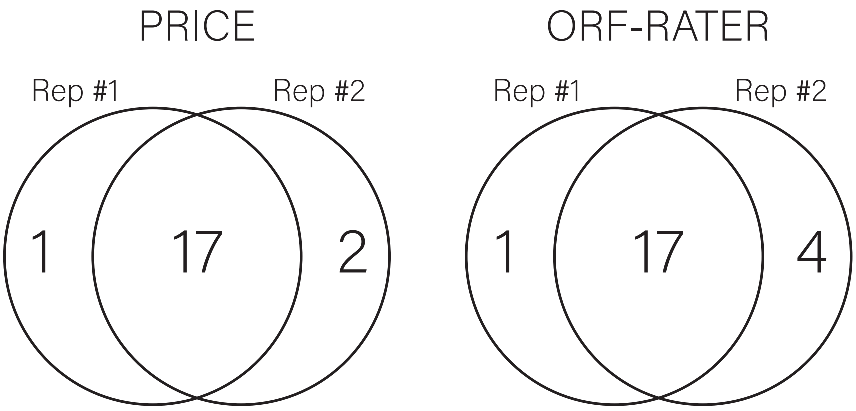


**Figure S14.** Reproducibility of viral ORFs predictions

Venn diagram summarizing the reproducibility of the ORF classifiers (PRICE and ORF-RATER) between our independent biological replicates.

### Figure S15

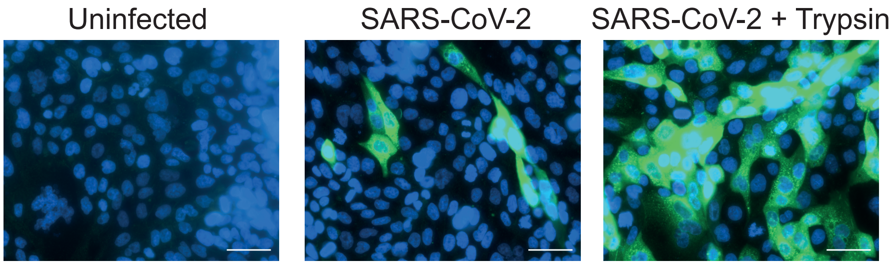


**Figure S15**. SARS-CoV-2 infection of Calu3 cells.
Calu3 cells were either left uninfected, infected with SARS-CoV-2, or infected with SARS-CoV-2 in the presence of trypsin. 12hpi the cells were fixed and stained with antisera against SARS-CoV-2 (green) and Dapi (blue).
Representative microscopy images are presented. Scale bars are 200μm.

Figure S16

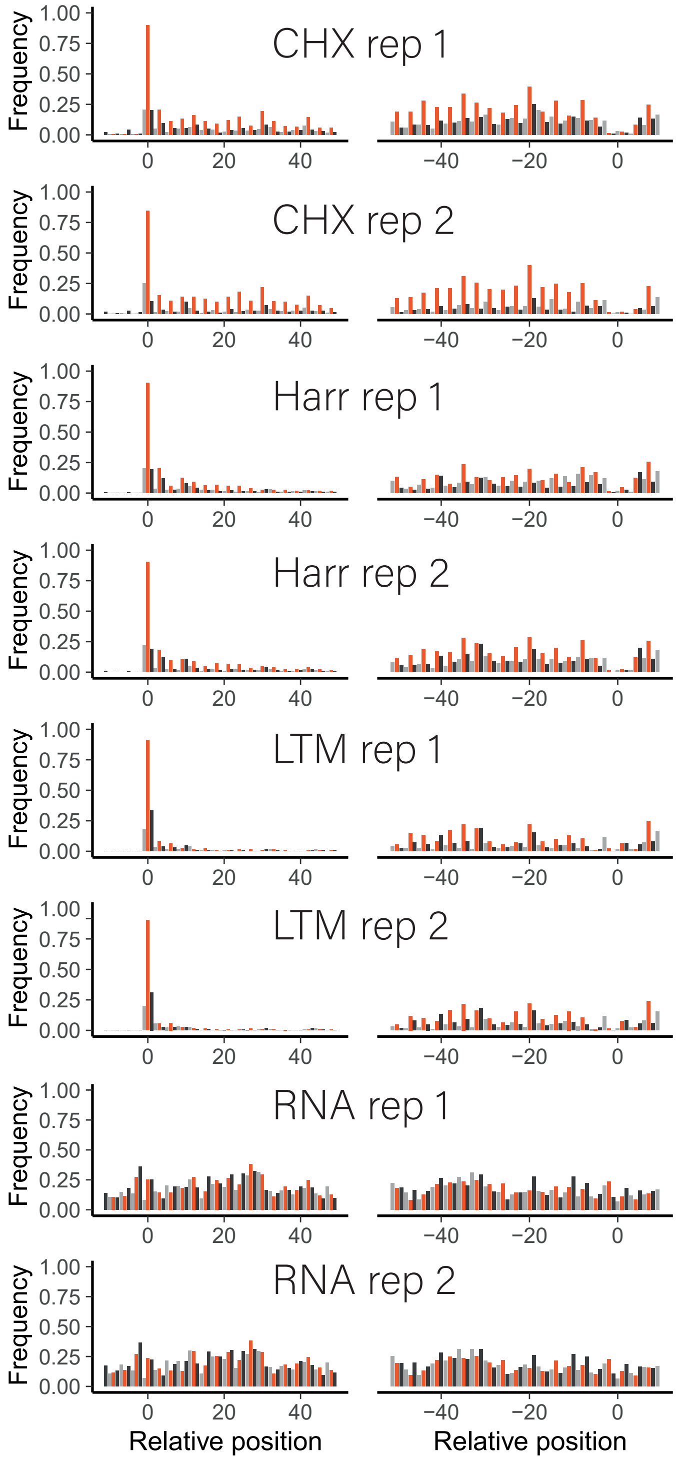


**Figure S16.** Footprint profiles of viral coding genes from SARS-CoV-2 infected Calu3 cells

Metagene analysis of read densities at the 5' and the 3' regions of viral protein coding genes as measured by the different ribosome profiling approaches and RNA-seq at 7hpi from two biological replicates. The X axis shows the nucleotide position relative to the start or the stop codons. The ribosome densities are shown with different colors indicating the three frames relative to the main ORF (red, frame 0; black, frame +1; grey, frame +2).

### Figure S17 - page 1

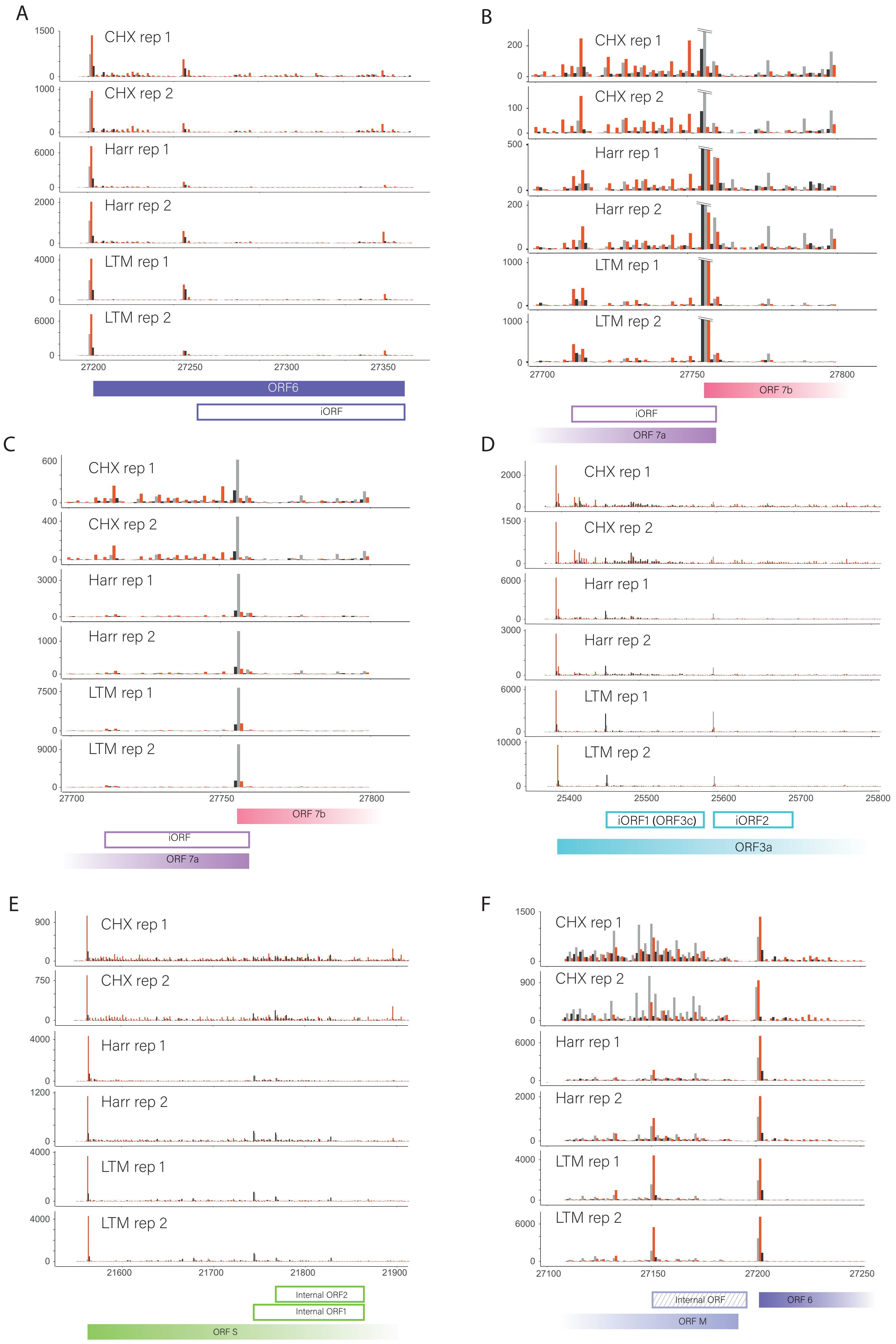

Figure S17 - page 2

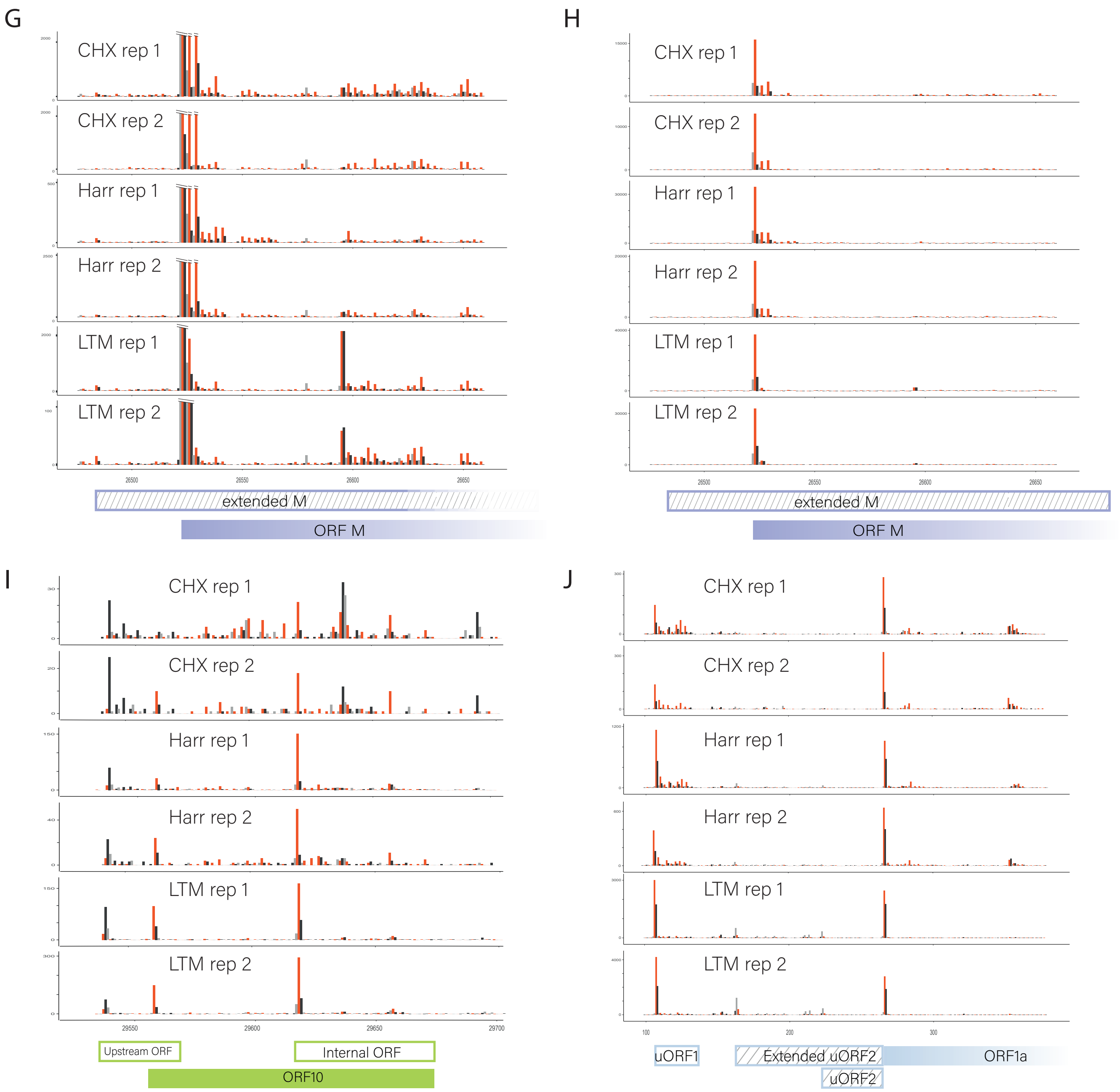


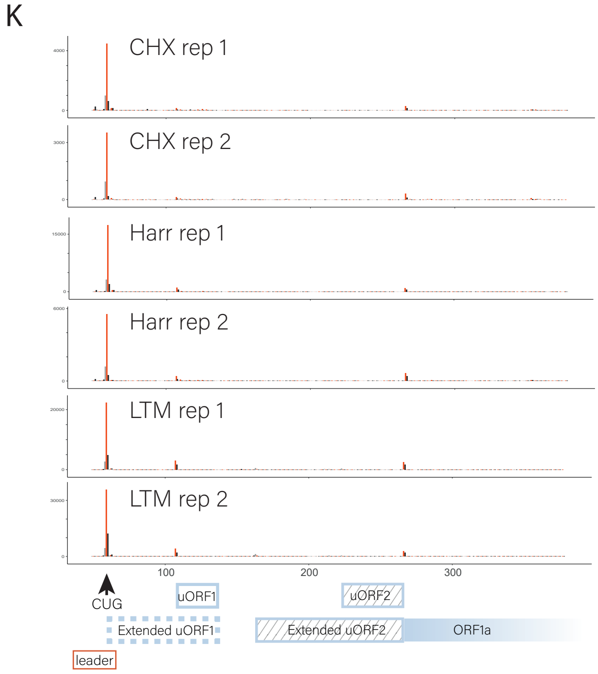


**Figure S17**. Footprint profiles reveal novel viral coding regions are also translated in infected Calu3 cells.
(A-K) Read densities over SARS-CoV-2 unannotated translated ORFs as measured by the different ribosome profiling approaches from both replicates at 7hpi. The ribosome densities are shown with different colors indicating the three frames relative to the main ORF in each figure (red, frame 0; black, frame +1; grey, frame +2). Filled and open rectangles indicate the canonical and novel ORFs, respectively. ORFs starting in near cognate start codon are labeled with stripes. (A) In frame internal initiation within ORF6 generating truncated product. (B and C) In frame internal initiation within ORF7a, (B) scale is adapted to present the internal ORF initiation (C) full scale presenting ORF7b initiation in true scale. (D) Out of frame internal initiation within ORF3a. (E) Out of frame internal initiation within ORF-S. (F) Out of frame internal initiation within ORF-M. (G-H) an extended version of ORF-M (G) scale is adapted to present the extended ORF (H) full scale presenting the dominance of the main initiation. (I) uORF that overlap ORF10 initiation and in frame internal initiation generating truncated ORF10 product (I) two uORFs embedded in ORF1ab 5'UTR (J) non-canonical CUG initiation upstream of the TRS-leader

Figure S18

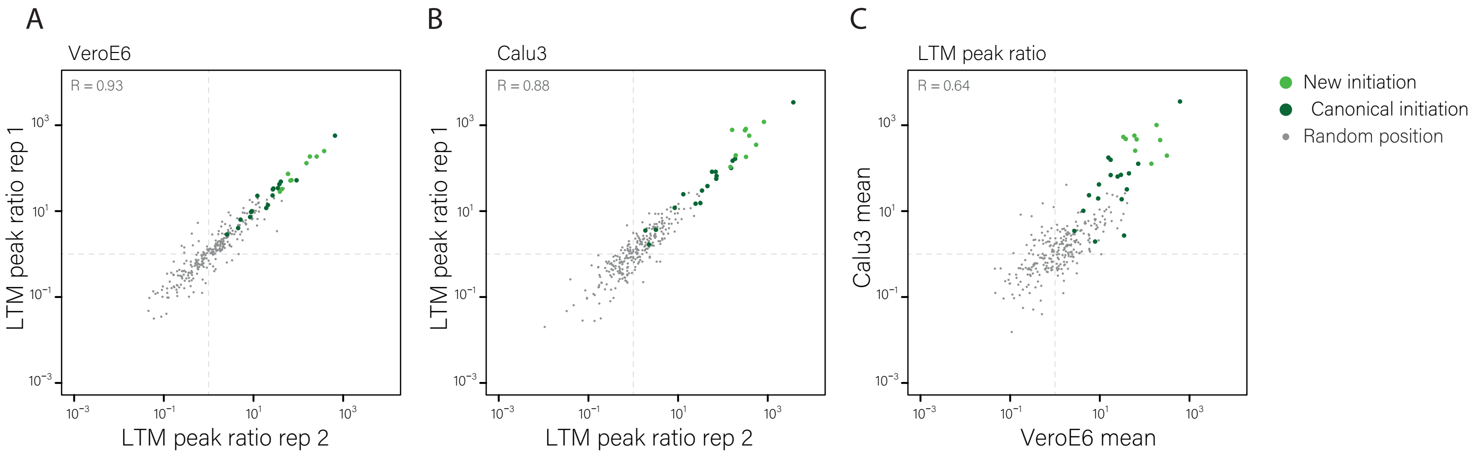


**Figure S18** LTM- induced ribosome accumulation at initiation sites are highly reproducible.

Scatter plot presenting the correlation of footprint densities at each initiation site relative to a 8bp window 3 nucleotides downstream of the initiation site in LTM treated samples between two biological replicates of infected Vero E6, 5hpi **(A)**, between two biological replicates of infected Calu3 cells, 7hpi **(B)** and between infected Vero E6 cells at 5hpi and Calu3 cells at 7hpi **(C)**. Canonical ORFs are marked in dark green and newly identified ORFs are marked in light green. As a control, the relative occupancy of random positions was calculated in the same way (grey). Dashed lines mark the equal ratio of one. Spearman’s R is presented.

Figure S19

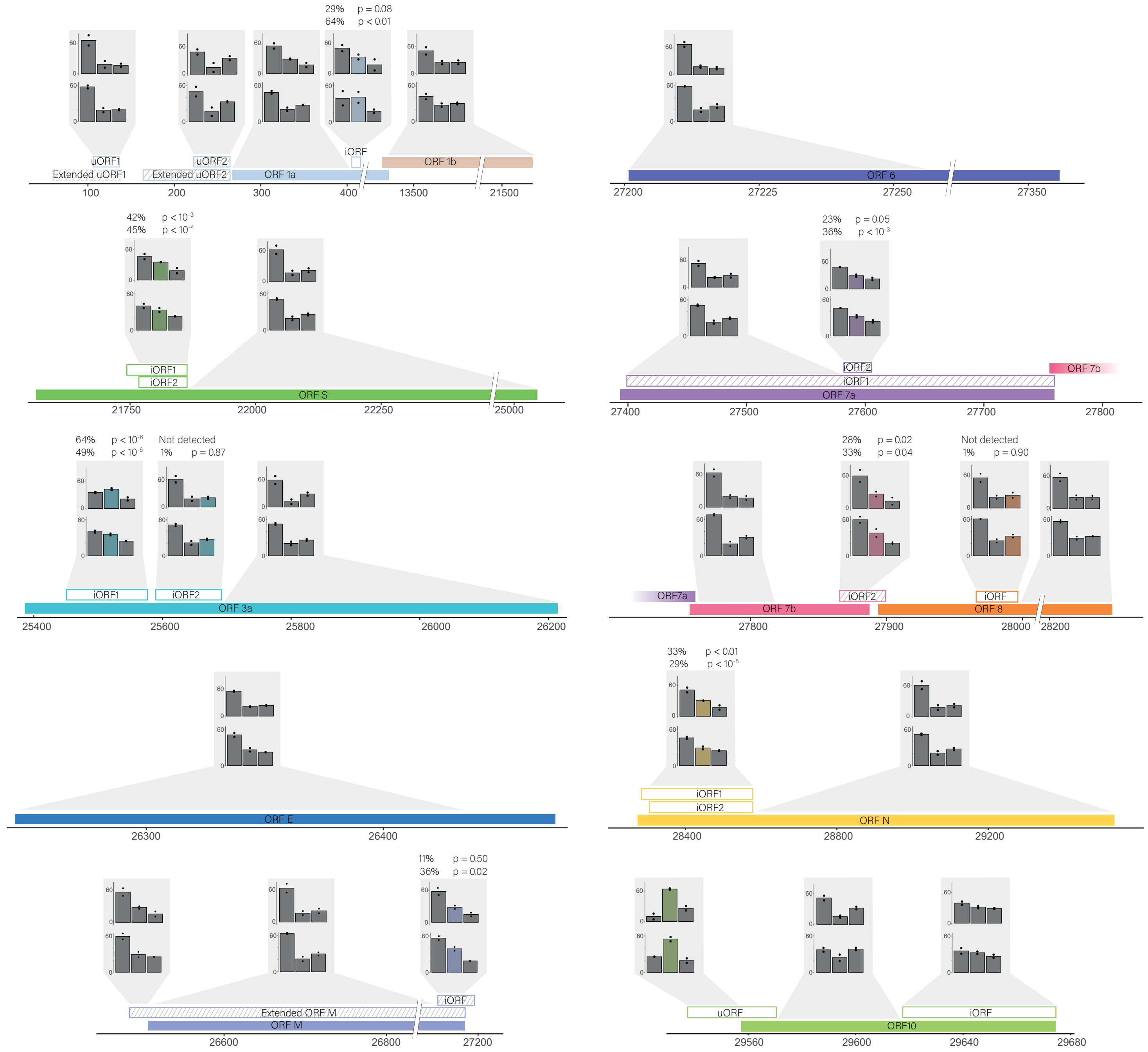


**Figure S19** Position of ribosome footprints relative to the reading frame in viral ORFs

The position of ribosome footprints relative to the reading frame in all canonical ORFs and novel ORFs excluding in-frame internal ORFs are presented for our measurements of infected Vero-E6 cells at 5hpi (lower panels) and infected Calu3 cells (upper panels). Filled and open rectangles indicate the canonical and novel ORFs, respectively. ORFs starting in near cognate start codons are labeled with stripes. The frame of the footprints is summed on each of the indicated regions and is presented relative to the frame of the canonical ORF in each of these loci. In all non-overlapping regions, beside ORF10 in Vero-E6 cells, clear enrichment to the translated frame is observed indicating active translation. For out-of-frame overlapping ORFs the bar of its frame is labeled by color and the percentage of the footprints that originate from the overlapping frame as was calculated from linear regression is presented together with the corresponding P-value for the contribution of the out-of-frame ORF to the frame distribution of the total reads in this region. In two out-of-frame overlapping ORFs, ORF3.iORF2 and ORF8.iORF the expression relative to the main ORF was low and did not lead to a significant shift in the translation signal. In all other ORFs there is a significant signal in the alternative frame indicating active translation.

Figure S20

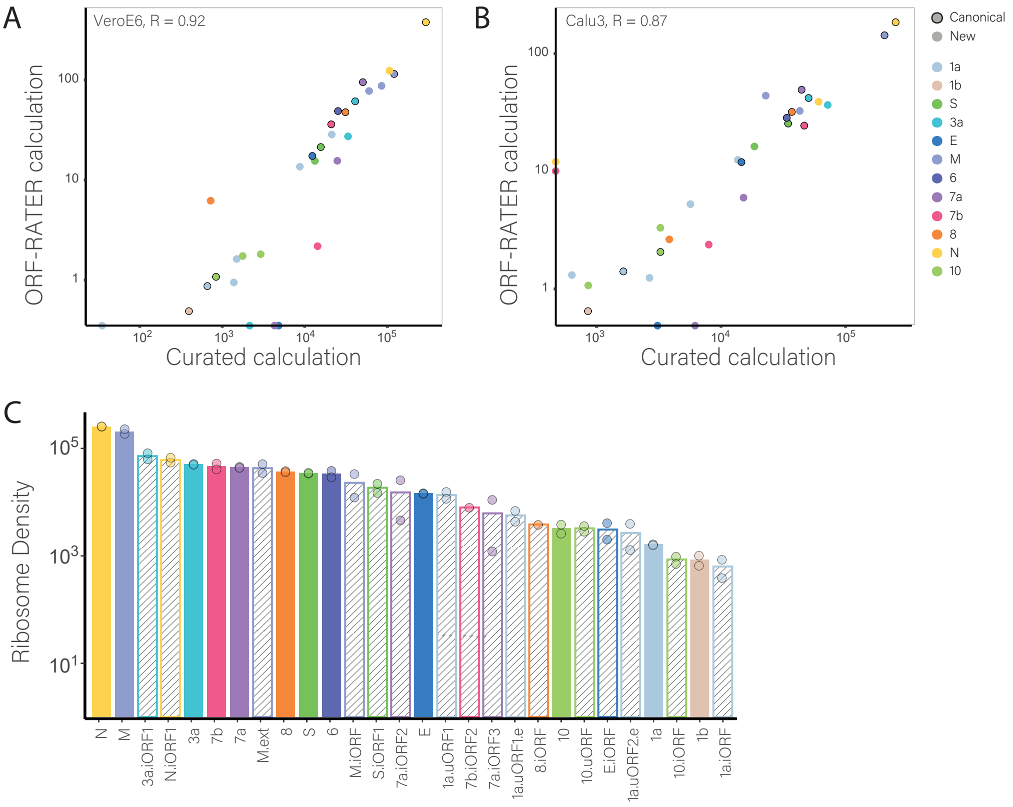


**Figure S20** Translation of viral ORFs

**(A-B)** Scatter plots presenting the correlation between translation levels as estimated by our curated quantification and as calculated by ORF-RATER for VeroE6 **(A)** and Calu3 cells **(B)**. Points representing canonical ORFs are outlined in black. Spearman's R is presented. **(C)** Viral ORF expression as calculated from ribosome densities of infected Calu3 cells. Data is plotted on a log scale to cover the wide range in expression. Solid fill represents canonical ORFs, and stripe fill represent novel ORFs that were annotated. Values were normalized to ORF length and sequencing depth. Points represent the values of each individual replicate.

Figure S21

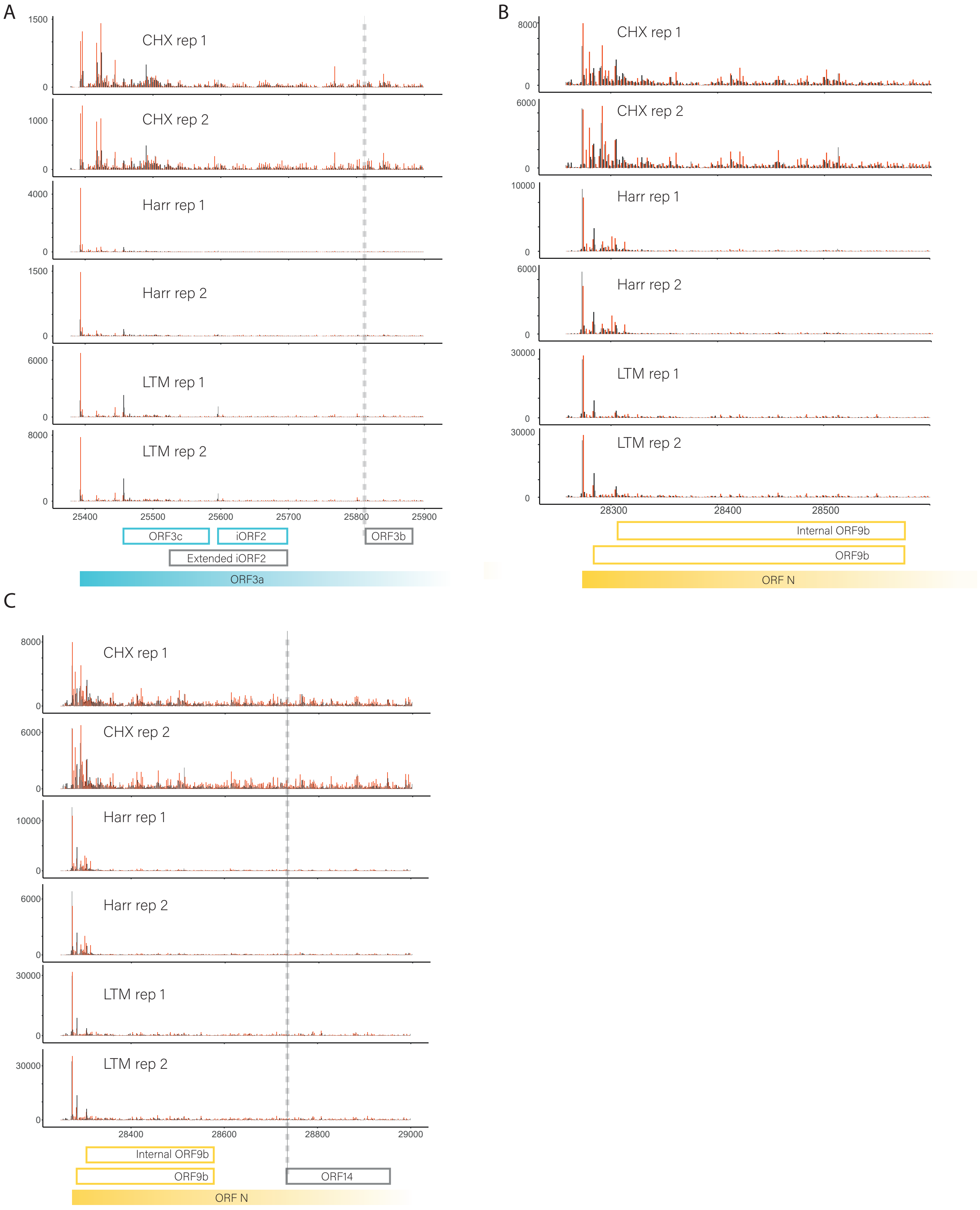


**Figure S21.** Internal ORFs encoded within ORF-N and ORF3a
(**A-C**) Read densities over ORF3a **(A)** and ORF-N **(B and C)** as measured by the different ribosome profiling approaches in two replicates in VeroE6 cells at 5hpi. The ribosome densities are shown with different colors indicating the three frames relative to the main ORF in each figure (red, frame 0; black, frame +1; grey, frame +2). Filled and open rectangles indicate the canonical and novel ORFs, respectively. ORF3b, extended iORF2 and ORF14 are marked based on the homology to SARS-CoV.

Figure S22

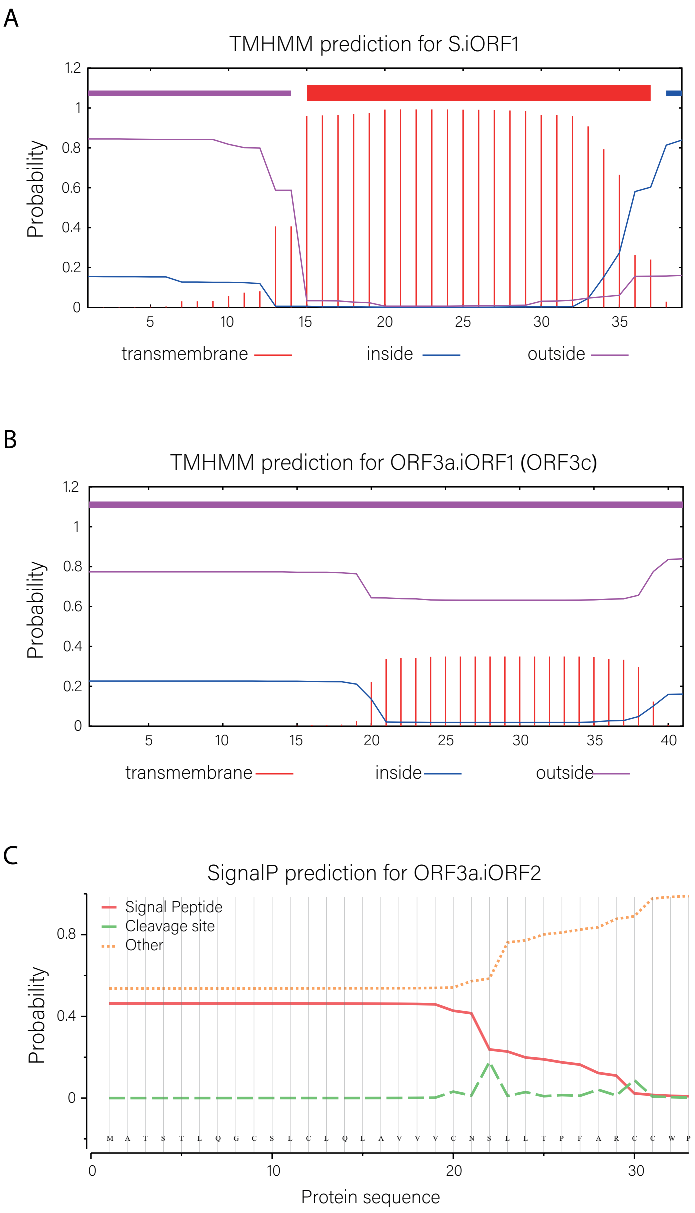


**Figure S22**. Transmembrane and signal peptide predictions
**(A)** Transmembrane region predicted in S.iORF1 using TMHMM 2.0 **(B)** Transmembrane region predicted in 3a.iORF1 (ORF3c) using TMHMM 2.0 **(C)** signal peptide prediction in 3a.iORF2 as predicted using SignalP 5.0.

Figure S23

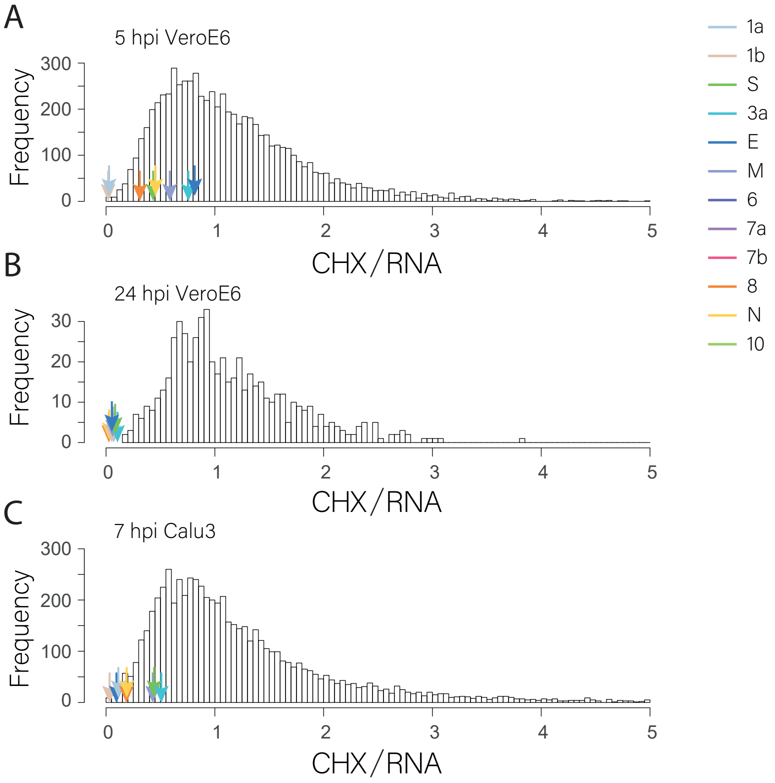


**Figure S23.** Comparison of host and viral translation efficiency

**(A -C)** Histograms showing translation efficiency of host genes calculated as the ratio between footprint densities and RNA abundance normalized to the median of each samples, at 5 hpi **(A)** and 24 hpi **(B)** in VeroE6 cells and at 7hpi in Calu3 cells **(C).** Arrows show translation efficiency for each viral gene. Histone transcripts are under-represented in poly-A selected RNAseq libraries, and were therefore excluded from these analyses.
